# The dynamic hypoosmotic response of *Vibrio cholerae* relies on the mechanosensitive channel MscS

**DOI:** 10.1101/2023.05.08.539864

**Authors:** Kristen Ramsey, Madolyn Britt, Joseph Maramba, Blake Ushijima, Elissa Moller, Andriy Anishkin, Claudia Häse, Sergei Sukharev

## Abstract

Like other intestinal bacteria, the facultative pathogen *Vibrio cholerae* adapts to a wide range of osmotic environments. Under drastic osmotic down-shifts, *Vibrio* avoids mechanical rupture by rapidly releasing excessive metabolites through mechanosensitive (MS) channels that belong to two major types, low-threshold MscS and high-threshold MscL. To investigate each channel’s individual contribution to *V. cholerae’s* osmotic permeability response, we generated individual *ΔmscS, ΔmscL*, and double *ΔmscL ΔmscS* mutants in *V. cholerae* O395 and characterized their tension-dependent activation in patch-clamp experiments, as well as their millisecond-scale osmolyte release kinetics using a stopped-flow light scattering technique. We additionally generated numerical models reflecting the kinetic competition of osmolyte release with water influx. Both mutants lacking MscS exhibited delayed osmolyte release kinetics and decreased osmotic survival rates compared to WT. The *ΔmscL* mutant showed comparable release kinetics to WT, but a higher osmotic survival, while Δ*mscS* had low survival, comparable to the double *ΔmscL* Δ*mscS* mutant. By analyzing release kinetics following rapid medium dilution, we illustrate the sequence of events and define the set of parameters that characterize discrete phases of the osmotic response. Osmotic survival rates are directly correlated to the extent and duration of cell swelling, the rate of osmolyte release and the onset time, and the completeness of the post-shock membrane resealing. Not only do the two channels interact functionally during the resealing phase, but there is also a compensatory up-regulation of MscS in the *ΔmscL* strain suggesting some transcriptional crosstalk. The data reveal the advantage of the low-threshold MscS channel in curbing tension surges, without which MscL becomes toxic, and the role of MscS in the proper termination of the osmotic permeability response in *Vibrio*.

## Introduction

All free-living microorganisms, both commensal and pathogenic, confront environmental challenges involving osmotic up- and down-shifts. This is especially relevant to intestinal microorganisms, which are primarily transferred between hosts through freshwater. These osmotic forces are strong; a typical accumulation of 0.2 M metabolites inside the cell above the medium osmolarity creates a total of 4.8 atm of pressure (Csonka and Hanson, 1991; Record et al., 1998). Thus, all free-living microorganisms must have defenses against drastic osmotic changes to survive (Cox et al., 2018; Kung et al., 2010). As bacterial membranes are permeable to water, their sole mechanism for evading osmotic rupture is by reducing the osmotic gradient. Bacterial mechanosensitive (MS) channels mediate osmotic forces by acting as osmolyte release valves. Under hyperosmotic conditions, bacteria will accumulate osmolytes compatible with their cellular functions (Wood, 2006; Yancey et al., 1982), such as K+, glycine betaine, and ectoine (Pflughoeft et al., 2003). In the case of hypoosmotic shock, a sudden dilution of external media leads to rapid swelling and pressure buildup within the cell. To survive and curb the pressure surge, the cell must expel the accumulated osmolytes at a rate that surpasses the water influx. Thus, survival under osmotic down-shock is a kinetic competition between these two fluxes (Bialecka-Fornal et al., 2015).

MS channels in bacteria were first discovered when the procedure of giant spheroplast preparation was adopted for direct patch-clamp recording (Martinac et al., 1987). Bacterial MS channels from the two separate MscS and MscL families (Sukharev et al., 1993) were then identified in *E. coli* (Levina et al., 1999; Sukharev et al., 1994). MscS, the small-conductance channel, opens at non-lytic tensions (5-7 mN/m) (Belyy et al., 2010; Sukharev, 2002) and exhibits adaptive behavior that includes inactivation (Akitake et al., 2005). Conversely, MscL, the larger-conductance emergency release valve (Sukharev et al., 1994), opens at near-lytic tensions (10-14 mN/m) (Belyy et al., 2010; Sukharev et al., 1999) and does not inactivate. MS channels have since been found in all free-living bacteria, both gram-negative and gram-positive (Balleza and Gomez-Lagunas, 2009). More attention was given to opportunistic pathogens such as *Pseudomonas aeruginosa* (Cetiner et al., 2017) and *V. cholerae* (Rowe et al., 2013), where these channels were electrophysiologically characterized in some detail. In addition to electrophysiology, the physiological roles of individual bacterial MS channels are typically researched through generation of knockout mutants lacking one or more channels. Their osmotic phenotypes are characterized in hypotonic shock survival experiments (Levina et al., 1999), and more recently the application of the stopped-flow light scattering technique has been developed to study fast kinetics of osmolyte release (Cetiner et al., 2017; Moller et al., 2023).

While *E. coli* is primarily found in normal flora of the gastrointestinal tract, *V. cholerae*, like many species of *Vibrio*, lives in brackish waters, frequently attached to crustacean shells (Nahar et al., 2011). *Vibrio cholerae*, a typical estuarial marine bacterium, becomes a deadly pathogen when it acquires and incorporates a complex of virulence genes horizontally passed through phages or plasmids. Being adaptable to both high-salinity and fresh water, toxicogenic strains of *V. cholerae* often cause seasonal outbreaks of cholera in areas where potable, clean water is unavailable (Emch et al., 2008). *V. cholerae* osmoadaptivity correlates with the presence of MscL, which is absent from many strictly marine members in the *Vibrio* genus (Booth and Blount, 2012). MscL appears to be a factor that allows *V. cholerae* to colonize and infect the human gut without rupturing (Keymer et al., 2007) during fresh-water transfer between individuals. Although *V. cholerae* pathogenesis is known to be dependent on the presence of various phage-associated virulence factors (Faruque et al., 1998), its easy transfer and propagation appear to be contingent upon the presence of both MscS and MscL.

To achieve a mechanistic understanding of *Vibrio’s* osmoregulation and environmental stability, in this work we generated the Δ*mscL* and Δ*mscL ΔmscS* mutants in *V. cholerae* O395 toxT:lacZ (treated as WT) and characterized them, along with a previously developed Δ*mscS* mutant (Hase, 2001). We performed osmotic viability testing on each MS channel deletion mutant, along with WT, to characterize the channels’ individual contributions and collective effects on osmoadaption. We fully exploit the stopped-flow light scattering technique and quantitative analysis of scattering traces supplemented with extensive kinetic modeling that separates the release process into distinct phases reflecting cell mechanics and permeability. The comparison of WT *V. cholerae* O395 with isogenic Δ*mscS*, and the double Δ*mscL ΔmscS* mutants revealed that MscS is critical for osmotic survival and that the remaining MscL does not functionally substitute for MscS. To single out MscS’s function, we independently generated a Δ*mscL* mutant and observed that the Δ*mscL* cells had a fast permeability onset, and the osmotic survival rate for the mutant was considerably higher than that of WT. This correlated with up-regulation of MscS as a compensation for the absent MscL verified by quantitative RT-PCR. This work demonstrates the critical role of MscS in the osmolyte release mechanism and resealing and suggests a yet unknown crosstalk between expression regulation of the *mscS* gene and the presence or absence of MscL.

## Materials and Methods

### Strains and Mutant Generation

The parental *V. cholerae* strain O395 toxT:lacZ (treated as WT), and the Δ*mscS* mutant previously generated from it were received from Dr. Claudia Häse. Using methods adapted from Ushijima et. al (Ushijima et al., 2016), we additionally generated a Δ*mscL* mutant and a double Δ*mscL ΔmscS* deletion mutant through homologous recombination of a deletion cassette into the relevant *V. cholerae* chromosome. The deletion cassette was delivered via suicide plasmid obtained from via tri-parental conjugation with *E. coli* (EC B3914).

### Osmotic Viability Assays

Osmotic viability was determined by subjecting cells to rapid down-shock and assaying for the survival percentage based on CFU’s/mL, normalized to a non-shock control. Standard Luria Bertani (LB) overnight bacterial cultures were diluted 1:100 into fresh LB media supplemented with NaCl to an osmolarity of 1200 mOsm. *V. cholerae* cells were then grown to early logarithmic phase (OD_600_ ∼0.25). To simulate environmental down-shock, 30 μL of the culture was rapidly pipetted into 5 mL’s of either 1200 mOsm LB (non-shock control) or a lower osmolarity LB (800, 600, 400, 350, 300, 250, 200, 150 mOsm). Media osmolarity was measured prior to shock by three independent readings on the Osmomat 3000 Freezing Point Osmometer (Gonotech). Inoculated culture was incubated at room temperature for 15 minutes, then 30 μL was further diluted into a fresh 5 mL of the corresponding osmolarity. Shocked bacterial cells were then plated in duplicate and all resulting colonies manually counted the next morning. To determine percent survival, each colony count was normalized to the non-shock control. The average survival rates and standard deviations were based on at least seven independent experiments. Two-tail two-sample t tests assuming unequal variances were performed to determine significance.

### Light-Scattering Stopped-Flow Experiments

To observe and characterize the osmolyte release kinetics of each MS channel down-shock, we employed the rapid-mixing osmolyte efflux stopped-flow technique with light scattering detection. The technique allows us to discern the characteristic times taken by sequential processes of bacterial cell swelling, channel opening, osmolyte and water release and finally either cell recovery or mechanical damage by the down-shock.

For down-shocks, overnight *V. cholerae* cultures were diluted 1:250 into fresh 1200 mOsm LB media and grown to an OD_600_ of 0.25, then briefly spun down to pellet and resuspended in the resulting supernatant to an OD_600_ of 2.0 to concentrate the cells. Once concentrated, *V. cholerae* cells were rapidly mixed with lower osmolarity LB media in a 1:10 ratio in a two-syringe stopped-flow machine (Bio-Logic SFM-2000) with an 80 μl optical mixing chamber (Figure S1). The stopped-flow machine was equipped with a modular spectrophotometer to measure the small-angle (5-13°) light scattering as a result of changes in cell size and refractive index (n) reflecting the concentration of internal non-aqueous components (“dry weight”) (Koch et al., 1996). Both cell size and n change due to water influx and subsequent osmolyte release when cells are rapidly shocked in lower osmolarity media.

Lower osmolarity LB media was made by diluting 1200 mOsm LB media with sterile deionized water to lowered osmolarities of 150, 300 mOsm, 450, 600, and 900 mOsm, which were used for shocks, in addition to deionized distilled water (ddH2O). The 1:10 ratio of cells in 1200 mOsm media to diluted media thus created final down-shock osmolarities of approximately 100, 250, 400, 500, 650, and 850 mOsm.

Light-scattering traces were collected over a period of 4 seconds, with sampling every 10 milliseconds, for a total of 8001 data points per shock. Cells were shocked sequentially into each diluted media with five technical replicates per trial and averaged. Down-shock trials were repeated with at least six biological replicates per clone. Readings of background scattering were done with the pure, cell-less supernatant medium. These were later subtracted from readings of the *V. cholerae* cellular shocks as a baseline.

Down-shock scattering traces were fit from the initialization of MS channels opening to the point where osmotic pressure lessens and membrane tension reaches below subthreshold level and channels close (typically occurring in 0.5 seconds). Fitting was performed on MATLAB, to the previously derived (Cetiner et al., 2017; Moller et al., 2023), simplified version Rayleigh-Gans equation, 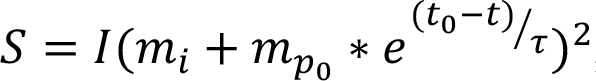, where S is the scattering and m_i_ and m_p_ are the fractions of impermeable and permeable osmolytes in masses, respectively. t_0_ designates the start of the fit, and τ is the rate of osmolyte release. t_0_ was chosen starting at the steepest negative slope (representative of MS channels opening) and fits then continued until this slope leveled off, when channels were observed to close. The analysis of light scattering traces included determination of the extent of initial cell swelling (ES), time to release onset (tro), time to the steepest point and the slope at the end of the trace.

### Spheroplast Preparation and patch-clamp

To generate giant spheroplasts for patch-clamp characterization, we adapted a previous *V. cholerae* spheroplast preparation technique used by our colleagues to make it more amenable to our selected O395 strain (Rowe, 2013). The techniques are based on the original procedure for giant spheroplast preparation from *E. coli* (Martinac, 1987). Briefly, cells were grown for 1.5 hours in MLB (250 mOsm) in the presence of 0.06 mg/mL cephalexin and 0.06 mg/mL carbenicillin, which prevents septation and causes them to grow as long chains of un-cleaved cells. Cells were then concentrated and resuspended into 1 M sucrose supplemented with 1.8 mg/mL BSA. The 15-min plasting reaction was developed at 110 rpm and 37C by adding 0.34 mg/mL lysozyme in the presence of 0.008 M EDTA, which breaks down the peptidoglycan layer and causes the long chains to collapse into giant spheres. The reaction is stopped with Mg2+ and spheroplasts are isolated via sedimentation through a sucrose gradient.

### RT-qPCR

The WT and Δ*mscL* strains were diluted 1:100 from overnight cultures into fresh LB and grown to early-log phase (OD_600_ of ∽0.25). Total RNA was extracted from cell cultures using the Monarch Total RNA Miniprep Kit (New England Biolabs) according to the manufacturer’s protocol. The concentration of RNA was measured with the NanoDrop OneC (Thermo Fisher Scientific) with an average yield of 185 ng/uL. RNA samples were then reverse transcribed into cDNA using the qScript cDNA Synthesis Kit (Quantabio), resulting in a final concentration of 10 ng/uL. The reverse transcription reaction was carried out at 22°C for 5 min, 42°C for 30 min, and 85°C for 5 min. RT-qPCR was carried out on a Roche LC480 using the PowerUp SYBR Green Master Mix (Thermo Fisher Scientific) according to the manufacturer’s protocol. Five technical replicates were run for four biological replicates of each strain with primers for both *mscS* and *adk*. The experimental cycling conditions were 50°C for 2 min, then 95°C for 2 min, followed by 40 cycles of 95°C for 15 s, 60°C for 15 s, and 72°C for 1 min. Data are presented as expression fold change using the 2^−ΔΔCT^ method relative to the consistently expressed housekeeping gene adenylate kinase, *adk* (Octavia et al 2013; Livak and Schmittgen, 2001). The error shown in Fig 8 is the standard deviation between replicates. A one-tail two-sample t test assuming unequal variances was performed to determine significance. Primer efficiency was determined using cDNA from both strains diluted in a 1:10 dilution scheme resulting in concentrations ranging from 1 ng/μL to 0.00001 ng/μL with six technical replicates for each primer. (Wong and Medrano, 2005). The cycling conditions were 95°C for 5 min followed by 40 cycles of 95°C for 5 s and 60°C for 30 s. A melt curve was run after the cycles for both the experimental conditions and the primer efficiency consisting of 95°C for 5 sec, 65°C for 1 min, and 97°C with a ramp rate of 0.11°C/s. Both the primer efficiency data and the experimental data were analyzed using Abs Quant/Fit Points to determine the Ct’s and Tm Calling to visualize melt curves and confirm that only one amplicon was present. The averages were taken between the technical replicate Ct’s. The primer efficiency was 2.076319 (105.35%) with a slope of -3.1515 for the MscS primers and 2.149385 (107.05%) with a slope of -3.0092 for the adk primers. Based on these efficiencies and the average Ct’s, the relative expression ratio was calculated using the equation 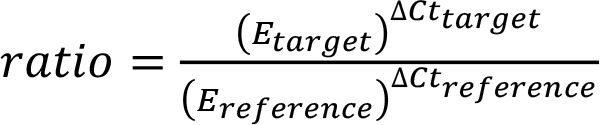 where E is the primer efficiency *E* = 10^−1/*slope*^. See Supplemental Figure S1 for the primers and primer efficiency plots.

### Kinetic modeling of the bacterial osmotic response

To integrate the information from this study and enhance the predictive power, we developed a detailed kinetic model of the tension-driven permeability response involving water influx, channel activation followed by osmolyte efflux and cell contraction due to water release and recoil of the cell wall. The model is presented as a system of interdependent Ordinary Differential Equations. The time course of the osmotic response is generated by numeric integration of the model using the COPASI software (Hoops et al., 2006; Mendes et al., 2009), an advanced simulator for biochemical networks that has been used previously to model osmoregulation in yeast (Spolaor et al., 2022).

A detailed description of all the variables, parameters, and equations of the model is provided in the Supplement. Briefly, the model has explicit variables such as cell volume and internal osmolyte concentrations classified as small and large permeable as well as impermeable, membrane tension, and open probabilities for channels, all changing with time (Fig. S4). The model also includes the possibility of membrane ‘crack’ formation when tension exceeds lytic threshold. It also has sets of explicit physical parameters and constants governing each transition, determined either from our own experimental measurements (e.g. expansion areas and gating energy for MscS and MscL, membrane permeability for water, starting refraction index of the medium, etc.), or from the literature (e.g. membrane elasticity and cell wall rigidity).

Some of the static parameters have to be adjusted for our specific system, for example because they might be only known for other bacterial species (e.g. cell wall rigidity), or because they might vary for every strain or the growth condition (e.g. the number of expressed/active MscS or MscL channels). COPASI enables automated iterative adjustment of the parameters within the expected range based on optimization of the fit between the predicted system behavior and experimental data. We used the predicted vs. experimental light scattering time course during the osmotic shock (stopped flow traces) to validate the model and adjust the needed parameters. To describe the complete time course, from the initial dilution of the cytoplasm with the same number of the osmolytes, and the following stage of the osmolyte loss, we used a more detailed version of the Rayleigh-Gans equation (Koch, 1961; Koch et al., 1996) that considers both cell volume and solute concentration changes in modeled osmotic response:

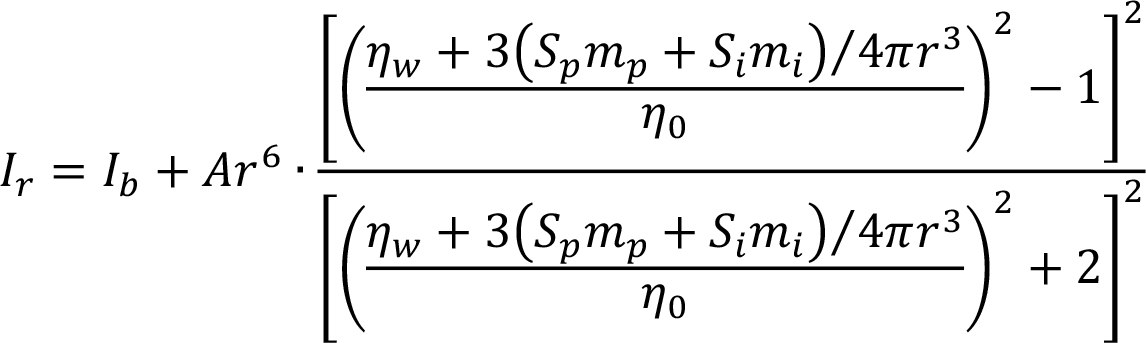

Here *I*_*r*_ is the intensity of the registered light, *I*_*b*_ is the background light component, *r* is the radius of the spherical equivalent of the bacterium, η_*w*_ is the refraction index of water, η_0_ is the refraction index of extracellular medium, *m*_*p*_ and *m*_*i*_ are the masses of the permeable and impermeable solutes respectively, *S*_*p*_ and *S*_*i*_ the scaling coefficients between their concentrations and contributions to the refraction index of the medium, and *A* is the scaling coefficient for the amplitude of the scattered light that combines components for the instrumental amplification, system geometry, angular dependence, light intensity, etc. The intracellular mass *m*_*p*_ of permeable osmolytes and their concentration are the main time-dependent variables (see Supplement for details).

## Results

### Generation of ΔmscL and double ΔmscS ΔmscS deletion mutants in V. cholerae 0395 and their primary characterization

V. cholerae strain O395 toxT::lacZ was taken as WT and used as the parent strain for each deletion mutant. In order to characterize each MS channel’s contributions to *V. cholerae*’s response to osmotic shock, we acquired the Δ*mscS* mutant (Hase, 2001) and generated deletion mutants for single Δ*mscL* as well as a double Δ*mscL* Δ*mscS*.

First, we confirmed each deletion mutant by subjecting the clones to PCR. For all mutants, screening primers resulted in amplicons that were smaller than the WT controls, indicating successful gene disruption (Figure 1). Next, successful MS channel deletions were functionally validated via patch clamp by testing for the presence or absence of channel characteristics that are unique to either MscS or MscL (Figure 2). As expected, WT exhibited two separate, distinct waves of channel activation under an increasing pressure ramp (Figure 2A). The initial channel activation at a lower threshold tension is MscS. As pressure increased and near-lytic tension was reached, a second wave can be seen, representing MscL opening. Conversely, in patch-clamp experiments on our Δ*mscS* clone, we only observed one single wave of activation (Figure 2B). As this activation wave was large and occurred at higher pressure, similar to that of MscL in WT *V. cholerae* O395, we can conclude that the only major channel activity present in Δ*mscS* is indeed MscL and that there are no obvious channels with MscS-like activity. Similarly, patch-clamp experiments run on our Δ*mscL* clone also showed a single wave of activation, this one at low pressures. The lower pressure wave is representative of MscS activation (Figure 2C). Interestingly, in double knockout spheroplasts we still observed some small-conductance and low-threshold channel activity (Figure 2D) that may represent the product of one of the remaining *mscS*-like genes such as the putative mechanosensitive channel protein given by accession ABQ22121.1 (BLAST: 292 aa, 37% identity to *E. coli* MscS).

**Figure 1.**
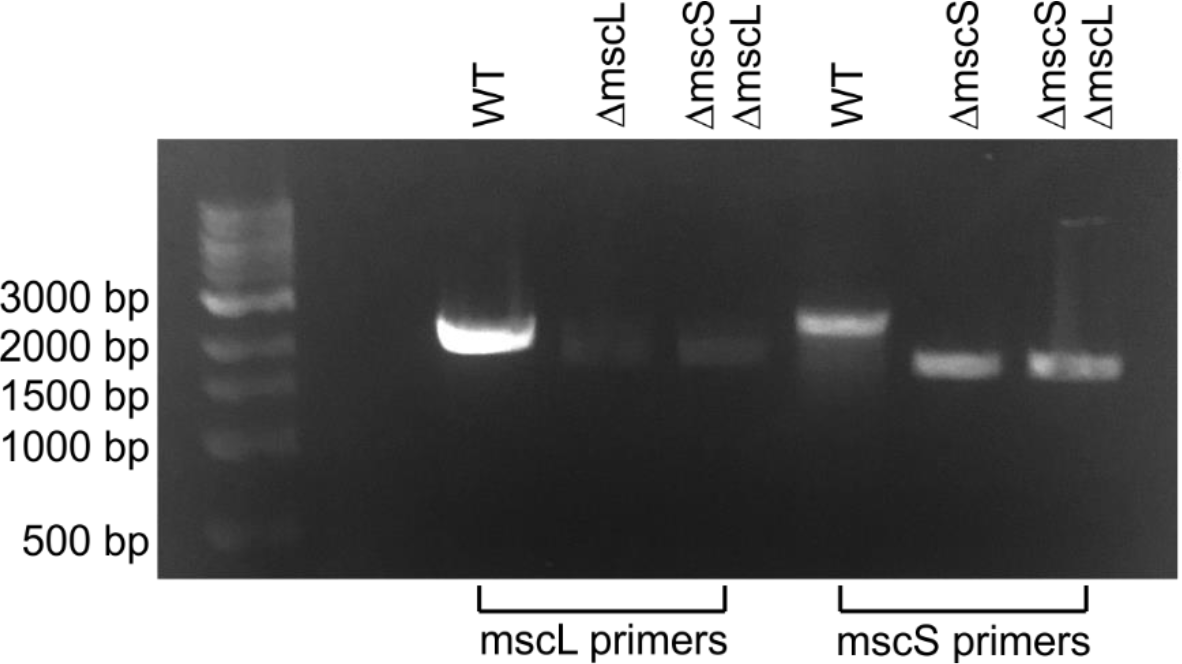
Generation of successful *ΔmscS, ΔmscL*, and *ΔmscS ΔmscL* knock-outs in VC O395. The gel shows the PCR bands generated with external diagnostic primers. Since the deletion cassettes are shorter than the genes, successful knockouts will produce amplicons that are shorter than their WT counterparts.

**Figure 2.**
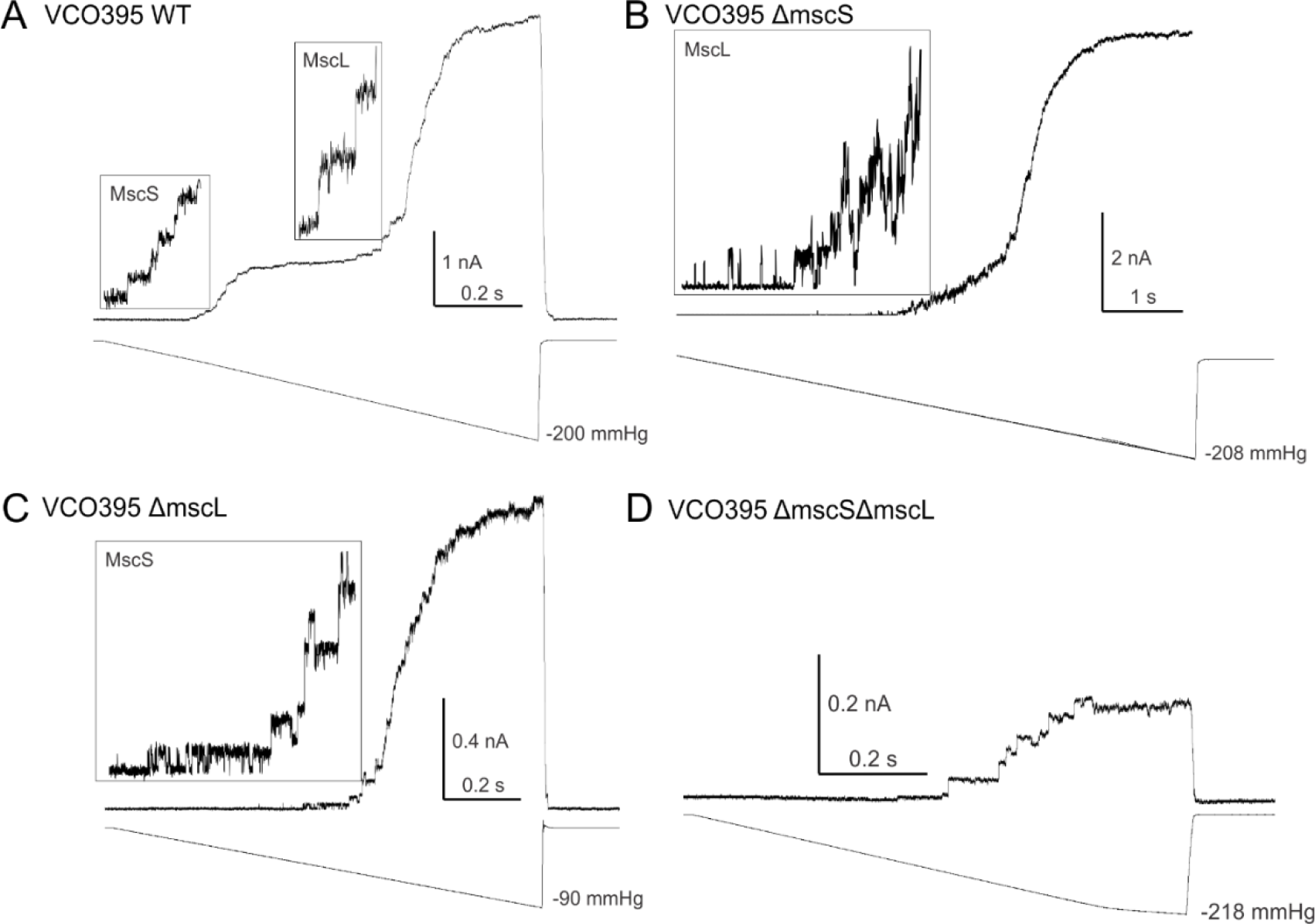
Electrophysiological characterization of WT and deletion strains by patch-clamp recordings from giant VC spheroplasts. (A) WT strain shows a characteristic ‘double wave’ generated by the populations of low-threshold (MscS) and high-threshold (MscL) channels. (B) The *ΔmscS* and (C) *ΔmscL* strains generate single waves reflecting the remaining type of channels. (D) The double *ΔmscS ΔmscL* mutant shows remnant channels without the characteristic waves of MscS and MscL.

### Characterization of V. cholerae O395 osmotic viability

To first understand the role of each MS channel to *V. cholerae* survival under hypoosmotic shock, we subjected WT *V. cholerae* and all MS channel KO mutants to osmotic viability assays. Surprisingly, these assays demonstrated a higher survival percentage under osmotic down-shock in Δ*mscL* than observed in WT *V. cholerae*. This increased survival occurred most prominently with large magnitudes of down-shocks (800 – 950 mOsm). The Δ*mscL* strain exhibited a survival rate below 50% only when down-shocked to an osmolarity of 200 mOsm, while WT *V. cholerae* reached 50% survival at shocks down to approximately 400-350 mOsm. Both Δ*mscS* and the double KO Δ*mscL ΔmscS* strains were observed to have a much lower tolerance for osmotic down-shock than WT, with both under 10% survival for shocks down to 400 mOsm. This result provides an unexpected prediction that MscL alone is not only insufficient to rescue the cells in this regime, but that all by itself it becomes toxic in the absence of MscS. This is consistent with the idea that MscS, with its distinctive ability to inactivate, helps terminate the massive permeability response by driving tension away from the MscL threshold (Akitake et al., 2005; Moller et al., 2023). At stronger shocks, however, MscL provides some advantage as a small number of Δ*mscS* colonies are found persisting in the shock plates.

### Characterization of V. cholerae O395 deletion mutants using stopped-flow

To understand the contributions each MS channel provides to the overall hypoosmotic permeability response, we performed rapid-dilution stopped-flow on each MS channel mutant to observe the kinetics of osmolyte release and correlate these kinetics with osmotic survival.

The principle of this method is based on the Rayleigh-Gans theory (Koch, 1961; Koch et al., 1996), where the forward scattering is proportional to the sixth power of the cell radius and to the square of the ratio of refractive indexes inside and outside the cell. The refractive index inside the cell, in turn, is directly proportional to the weight fraction of non-aqueous compounds (Moller et al., 2023). Figure 4 depicts the typical scattering kinetics that result from fast (8 ms) mixing of a suspension of WT *V. cholerae* O395 pre-grown in 1200 mOsm LB into a 400 mOsm medium. The trace exhibits an ∼ 70 ms-long nearly linear decline in scattering representing initial swelling of the cell resulting in modest cytoplasm dilution leading to a drop of refractive index **n**. Swelling generates tension in the inner membrane which activates MS channels. The region of the curve with the steepest slope is fitted with the simplified equation 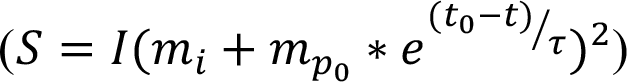 involving exponential release kinetics in order to extract the time constant. The scattering level at the end of the trace reflects the fraction of remaining (channel-impermeable) intracellular osmolytes contributing to the refractive index. This end level relative to the initial level also represents the fraction of permeable osmolytes that have exited the cell.

Vertical and horizontal lines in Figure 4 separate the scattering trace into distinct segments, beginning with the downward swelling phase, after which the trace shows a ‘break’, marked as time to release onset (tro), which is the minimum of the second derivative. The trace then enters the release phase by increasing its downward slope and passing through the steepest point. The minimum of the first derivative delineates the time to the steepest point (tsp). The curve then gradually flattens signifying the end of release and membrane resealing. The parameters of the release phase such as extent of swelling (ES), fraction of permeable osmolytes (FP), and slope at the end are quantifiable with the aid of calibration curves calculated according the Rayleigh-Gans theory ((Koch et al., 1996), supplement). This information can be used to calculate to what degree a cell increases its volume before tension in the inner membrane reaches the threshold and the MS channels open, how quickly they release excessive osmolytes, and what fraction of internal non-aqueous content leaves the cell. The slope at the end of the trace indicates the residual permeability and the quality of resealing.

Using these parameters, Figure 5 compares side-by side traces recorded in knockout strains versus the WT O395 strain. With all cultures, the “no shock” control traces where the cells are rapidly mixed with the 1200 mOsm ‘supernatant’ growth medium were flat. A qualitative observation between representative Δ*mscL* and WT scattering traces shows no sizable differences in the rates of osmolyte release throughout all magnitudes of down-shock (Figure 5A). Note that in the Δ*mscL* mutant, MscS is the dominant channel. We also observe similar trends in the end level reflecting the fraction of permeable osmolytes for the majority of down-shocks between WT and the Δ*mscL* mutant. For most shocks, by the 150 ms time point the scattering level is stabilized indicating that the release phase is over and tension has dropped to the threshold level for all MS channels in the population. However, two interesting differences can be seen at shocks to 900 mOsm and 400 mOsm. At down-shocks to 900 mOsm, WT appears to release less osmolytes overall than the Δ*mscL* clone, while the reverse is true at 400 mOsm (Figure 3A).

**Figure 3.**
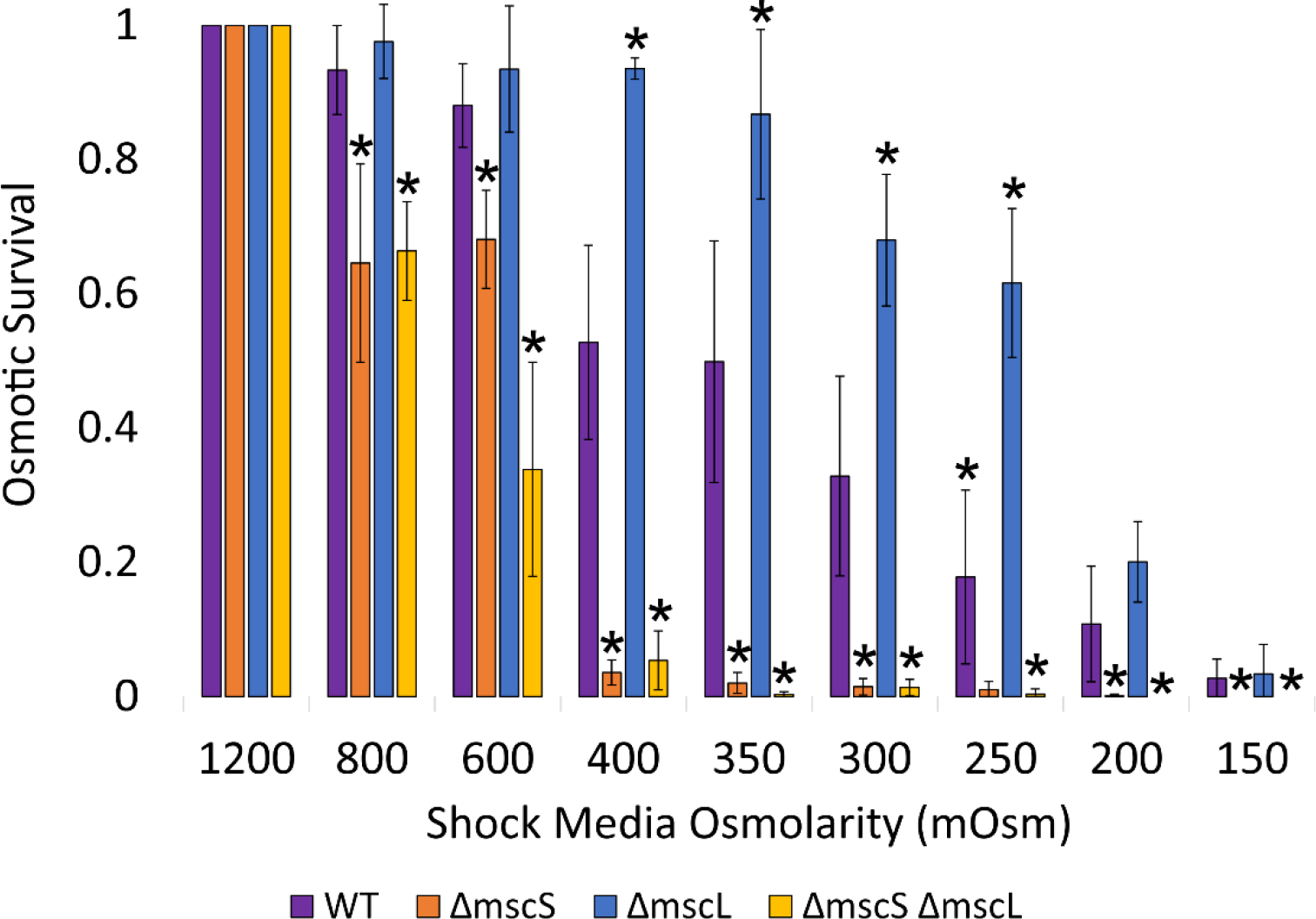
Osmotic viability of WT and MS channel deletion mutants. All samples were plated after osmotic shock from 1200 mOsm to the value indicated on the x-axis and survival measured by CFU’s/mL. Error bars represent standard deviation of each averaged sample (n=5-15). Significant values (< 0.05) are marked with an asterisk for values lower than WT, a pound sign for values higher than WT, or a plus sign for values that are higher compared to *ΔmscS*. The p-values are listed in table S1 and S2.

Converse to the response of the Δ*mscL* mutant, we observed that under osmotic down-shock, the Δ*mscS* mutant showed a delayed onset of downward deflection reflecting delayed channel openings with apparently higher tension threshold (MscL is the dominant channel) and slightly slower osmolyte release rates compared to WT *V. cholerae* O395. At intermediate magnitudes of shocks, the Δ*mscS* clone also appeared to expel a higher percentage of osmolytes than WT, as judged by the difference in scattering at comparable levels of shock. The faster and slightly deeper release in WT well correlates with its much higher survival rate especially at intermediate shocks relative to Δ*mscS* (Figure 3).

Upon deletion of both major release valves, the Δ*mscL ΔmscS* mutant exhibits a qualitative change in the scattering traces (Figure 5C). We should keep in mind that in this strain several minor MscS-like channels can still be expressed (Figure 2D) and may contribute to the release rate, but cannot impart high osmotic viability (Figure 3). We found that the double Δ*mscL*Δ*mscS* knockout strain exhibits a dramatically slower and more gradual release process. The traces do not show a typical sharp ‘break point’ as there is no sufficient population of channels that would synchronously activate at a given tension. The smooth break of the curve at high shock magnitudes instead reflects the formation of non-specific ‘cracks’ in the membrane. WT traces clearly indicate that at intermediate shocks the release process is complete by 150-200 ms. The analysis of entire 4 s traces of the double *ΔmscL ΔmscS* strain shows that at intermediate shocks osmolyte release does not stabilize until closer to 1.5 seconds following mixing. At high magnitudes of down-shock, to 500 mOsm and lower, the osmolyte release never appears to stabilize (Figure 5C). This strongly suggests that the release in the Δ*mscL ΔmscS* strain takes place through non-specific membrane ‘cracks’ which do not close or reseal. This pattern correlates with a drastic loss of viability (Figure 3).

Figure 6 directly compares traces recorded under identical 1200 to 400 mOsm downshifts in all four strains. We see that WT O395 cells are the most fit—their pre-release swelling is small, their release phase is fast (∼30 ms), they efficiently expel 11% of internal osmolytes, and they perfectly reseal. The Δ*mscL* strain carrying active MscS responds similarly quickly and stabilizes well after the shock even exhibiting a small upward drift likely reflecting slow cell shrinkage after osmolyte and water release possibly assisted by the cell wall recoil. The absence of the large channel, however, apparently changes the repertoire of permeable osmolytes and thus the fraction released at the end is smaller. The Δ*mscS* strain carrying active MscL shows a delayed activation, larger pre-release swelling and a prominent downward creep at the end of the trace indicating continuing release. The most ‘fragile’ double knockout Δ*mscS* Δ*mscL* strain shows the highest extent of pre-release swelling, which appears to be the major membrane-damaging factor. Correspondingly, the release through non-specific membrane cracks is slower, larger in magnitude, and never stops within the time scale of seconds.

All stopped-flow traces were fitted using MATLAB (Figure 4) to extract the rate of release, time to steepest point and the fraction of released osmolytes. The averages and standard deviations are plotted against the magnitude of osmotic shock and compared across all *V. cholerae* O395 WT and MS channel knockout strains in Figure 7. We found that there was no statistical difference between the release rates of Δ*mscL* mutant and WT *V. cholerae* at intermediate down-shocks. As noted in Figure 6, we see the largest difference between all mutants at intermediate magnitudes of down-shock, such as shocks to 400 mOsm. As expected, the double MS channel knockout, Δ*mscL* Δ*mscS*, had the slowest rates and a very gradual release consistent with a very different nature of permeation sites, which are likely tension-generated non-specific cracks rather than mechanosensitive channels.

**Figure 4.**
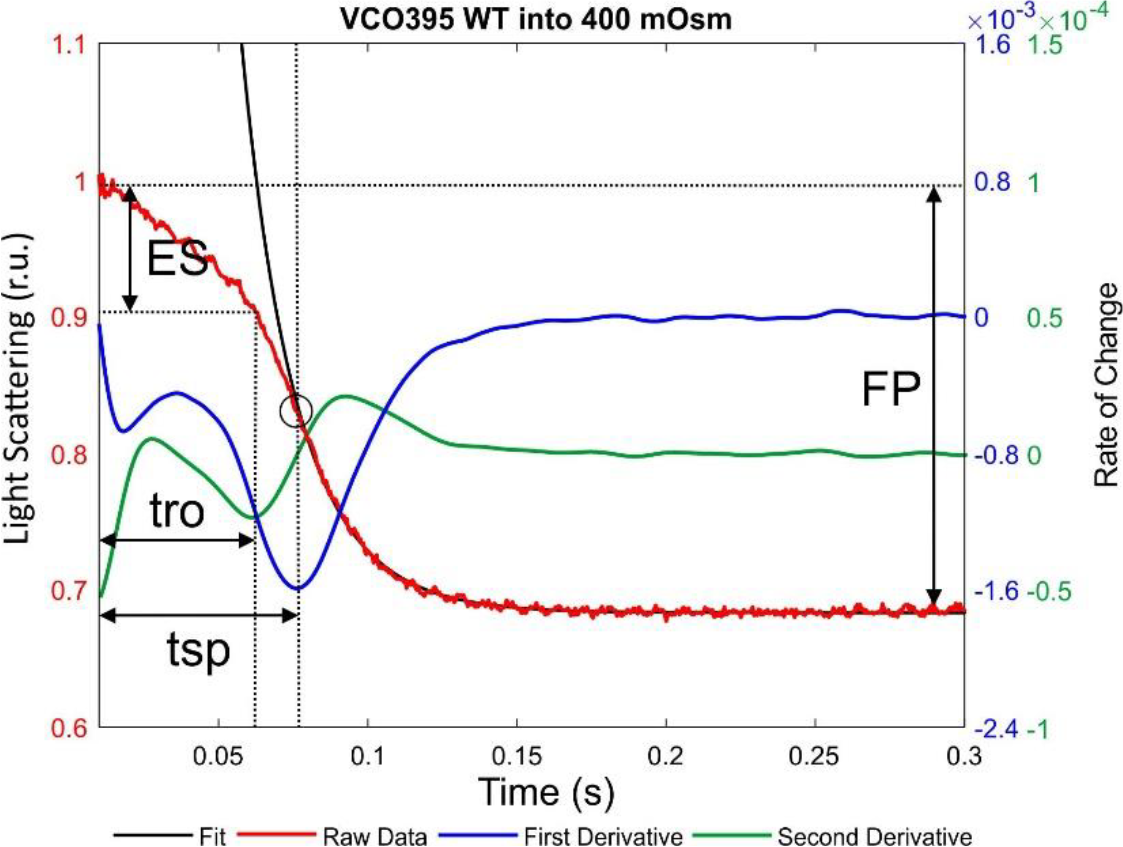
Representative scattering trace recorded in WT *V. cholerae* O395 that has been separated into distinct segments using derivative minima and fitting. The parameters such as time to release onset (tro, the minimum of the second derivative), extent of swelling (ES), time to the steepest point (tsp), fraction of permeable osmolytes (FP), and slope at the end are easily quantifiable via the first (blue) and second (green) derivatives. The rate of release is extracted from fitting the release phase with the simplified Rayleigh-Gans equation (black line).

**Figure 5.**
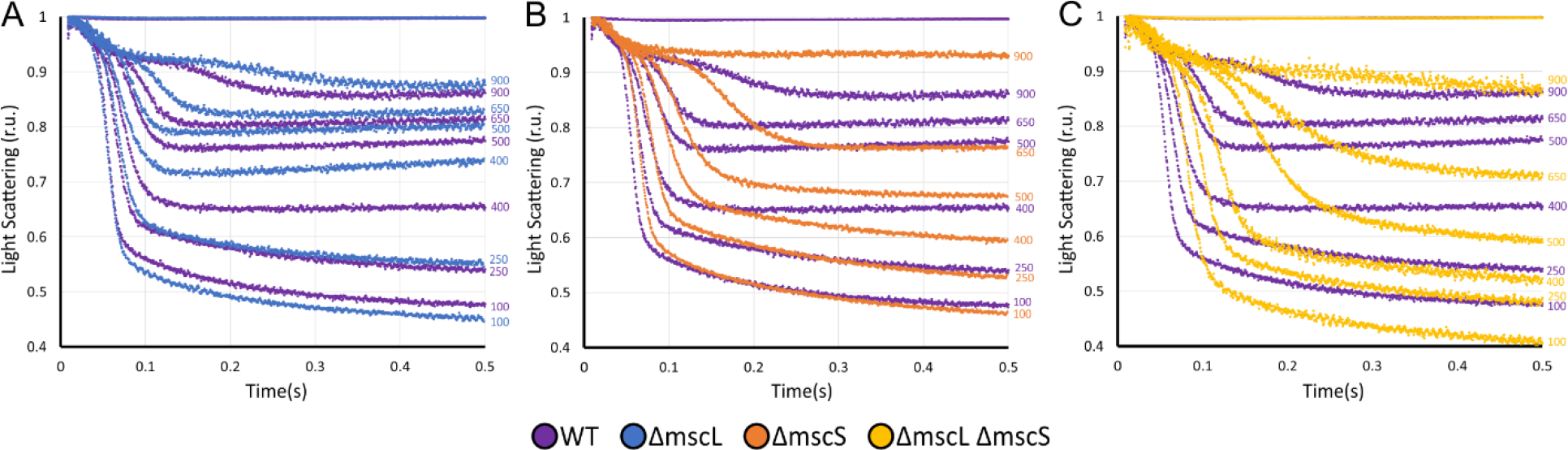
Osmolyte release kinetics recorded with light-scattering stopped-flow. The traces recorded in WT *V. cholerae* are shown in purple. Final osmolarities are indicated on the right. (A) WT cells versus Δ*mscL* mutant (blue). (B) WT versus Δ*mscS* mutant (orange). (C) WT versus Δ*mscL*Δ*mscS*. Note that the presence of high-threshold MscL as a dominant channel in the Δ*mscS* strain results in a slightly delayed onset of release and a stronger incline at the right end of the trace signifying incomplete resealing. All averages and standard deviations of kinetic parameters were results of 4-5 independent stopped-flow trials.

**Figure 6.**
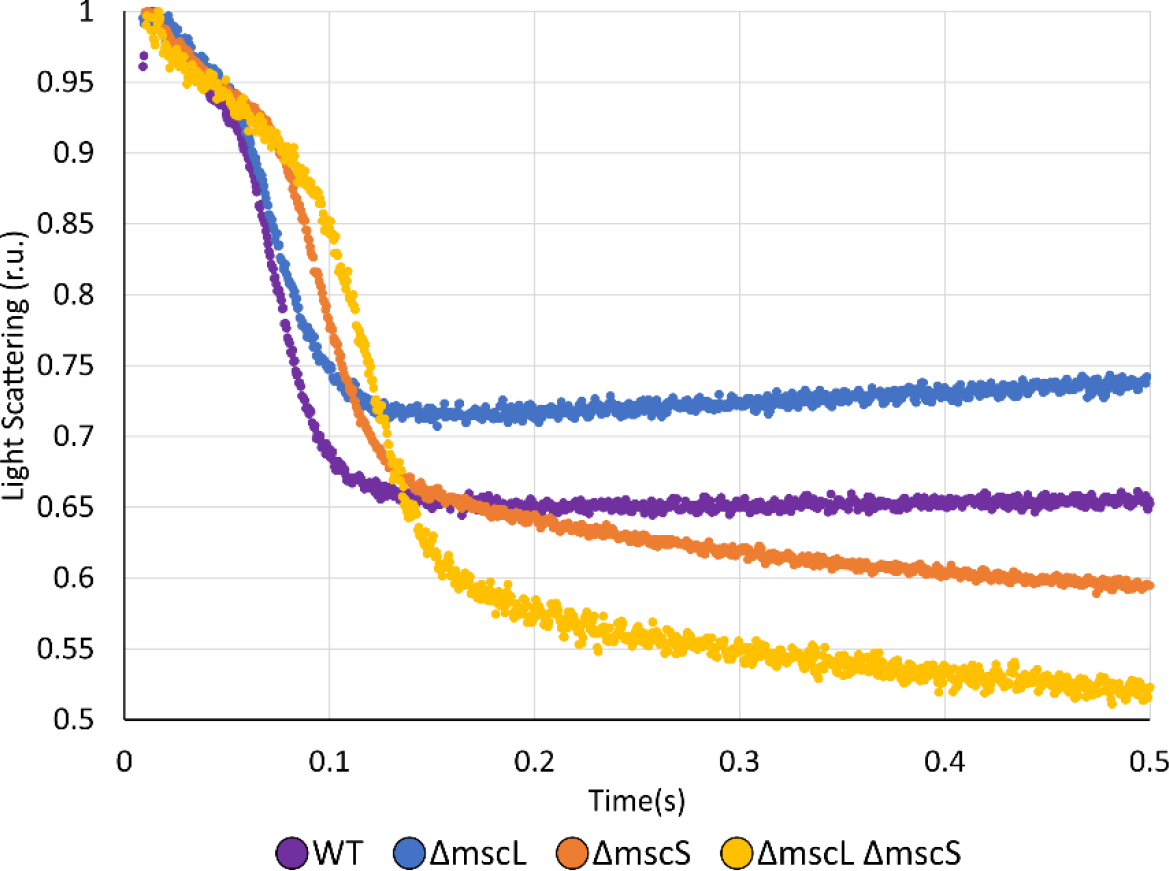
Comparison of light-scattering stopped-flow traces for different strains at the same osmolarity. Comparison of traces recorded under the same 1200 to 400 mOsm down-shock in WT *V. cholerae* cells (purple), Δ*mscL* (blue), Δ*mscS* (orange) and the double knockout Δ*mscS* Δ*mscL* strain (yellow). Note negative incline of traces signifying continuing leakage from Δ*mscS* and double knockout cells.

**Figure 7.**
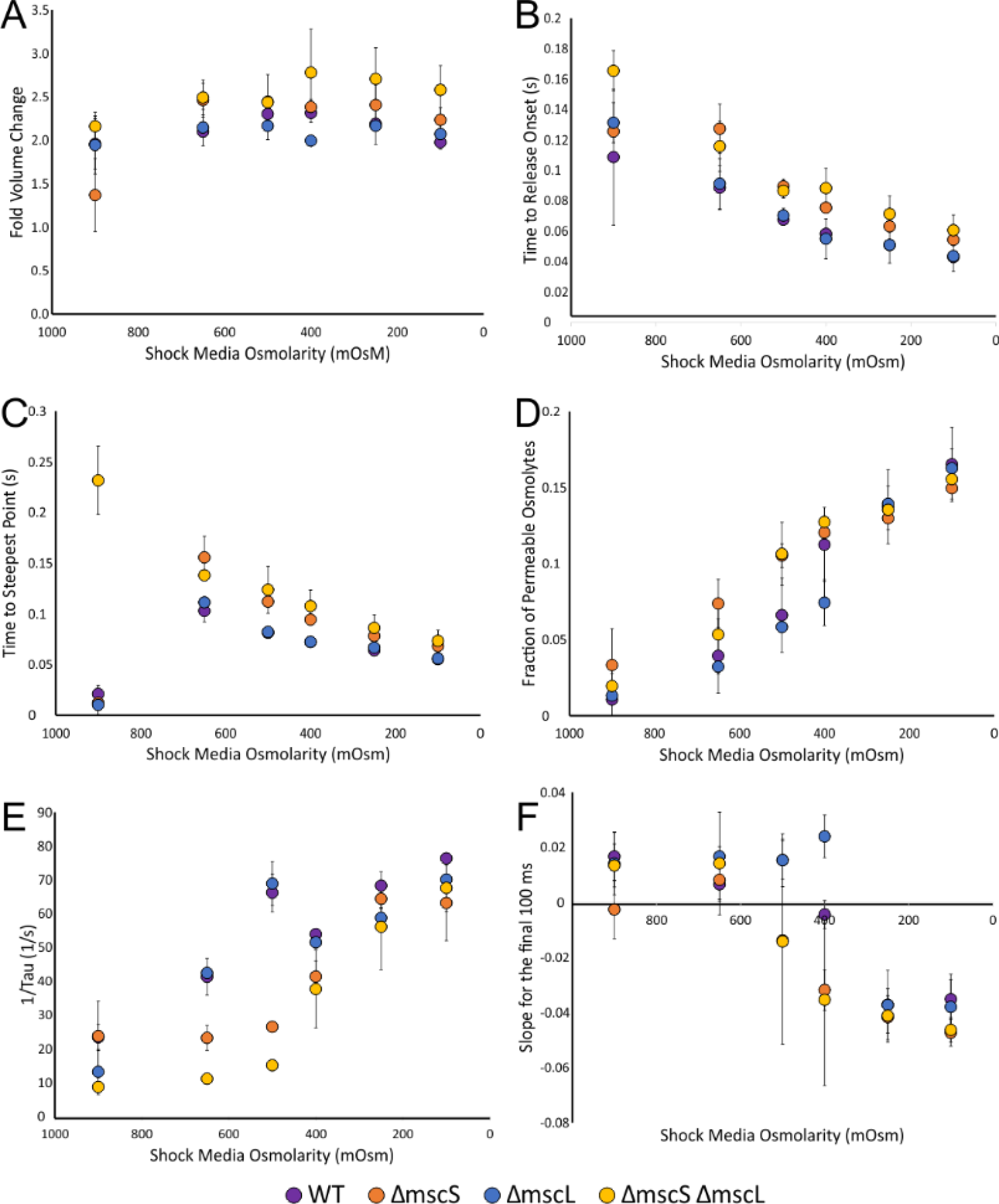
Kinetic parameters extracted from the stopped-flow traces recorded with *V. cholerae* O395 and the MS channel deletion mutants. Stopped-flow traces were fit to a modified Rayleigh-Gans equation from initialization of MS channel openings to the point of channel closure as described in Figure 2. These fits were averaged (n=4-5) and used to determine the osmolyte release rate under down-shocks from 1200 mOsm LB to the media osmolarity indicated on the x-axis. Error bars represent standard deviation. (A) The volume fold change as the cell swells in response to hypoosmotic shock. (B) Time to release onset corresponds to the time spent swelling until sufficient turgor pressure is achieved to activate channels. (C) Time to steepest point is the time at which the fastest scatter change occurred for each level of down-shock, indicating the time until maximum channel activity. (D) Fraction of permeable osmolytes released from *V. cholerae* O395. (E) The rate of osmolyte release. (F) The slope for the final 100 ms is a measure of membrane resealing. A positive slope is an indicator of good resealing, whereas a negative slope indicates the imperfect resealing that results in leaks through non-inactivating channels or nonspecific cracks.

We also examined the derivative of scattering changes for the point of steepest negative slope and time at which this occurred (Figure 4). This parameter suggested that at intermediate shocks (400 mOsm), the Δ*mscL* mutant likely has fewer active channels on average than other mutants. Yet, both WT and the Δ*mscL* mutant have comparably short times to the point of steepest slope at a majority of down-shocks (Figure 7C). The Δ*mscS* and double knockout Δ*mscL ΔmscS* mutants, however, are both significantly slower than WT and the Δ*mscL* strains to maximum channel activation.

The fractions of osmolytes that permeated through opened MS channels upon down-shock were also determined through the same fitting equation to stopped-flow data as release rates. At moderate shocks to 650 – 400 mOsm, the Δ*mscS* and Δ*mscL ΔmscS* mutants consistently released a higher fraction of permeable osmolytes than the Δ*mscL* and WT strains. This shows that MscS provides the most ‘economical’ regime of turgor adjustment. The two MscS-lacking mutants exhibited relatively similar percentages of permeable osmolytes at comparable shock levels to each other overall; however, as we saw from shocks to 400 mOsm, the double Δ*mscL ΔmscS* knockout clone continues to lose osmolytes until the end of the traces while the Δ*mscS* clone does not (Figure 7F). At large shocks from to 1200 mOsm down to 250 or 100 mOsm, which was always associated with higher death rate across the strains, *V. cholerae* O395 WT and MS channel knockout clones exhibited a similar trend (Figure 7D) showing a relatively steady increase in permeable osmolyte fraction. Again, this is likely due to the large contribution of leaking osmolytes from ‘cracks’ in damaged membranes, as this is a lethal shock. As predicted based on stopped-flow data, both Δ*mscS* and the double deletion Δ*mscL ΔmscS* strains were observed to have a much lower tolerance for osmotic down-shock than that of WT (Figure 3).

### MscS is overexpressed in the absence of MscL

To investigate the unexpected observations that the Δ*mscL* mutant both out-survives WT (Figure 3) and exhibits fast rates of osmolyte release (Figure 7E), we performed qRT-PCR to determine the expression level of native channels in Δ*mscL* compared to WT. We discovered that the MscS population in Δ*mscL* was upregulated approximately by 1.5 times (Figure 8), indicating that MscS is overexpressed in the absence of MscL.

**Figure 8.**
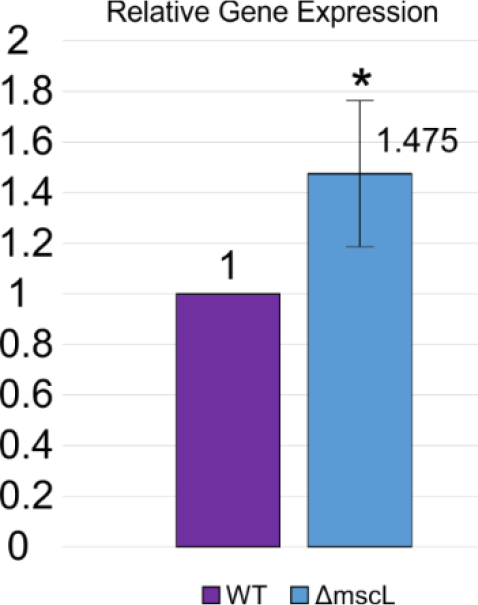
Native expression of MscS in WT and Δ*mscL*. qRT-PCR was performed to verify the overexpression of MscS observed in the electrophysiological characterization of the Δ*mscL* strain. Six independent experiments gave a ratio of 1.475 ± 0.289. Statistically significant (P = 0.02) expression level when compared to the WT strain is represented with an asterisk.

### Kinetic modeling of the cell-rescuing osmotic permeability response

The kinetics of osmolyte release measured using the stopped-flow technique now provides grounds for kinetic modeling of the *Vibrio* osmotic permeability response and the rescuing mechanism. The model is programmed in the biochemical kinetics suite *COPASI* (Hoops et al., 2006; Mendes et al., 2009) and described in the Supplement. It presents the bacterial cell as a compartment delineated by the cytoplasmic membrane surrounded by a porous and elastic cell wall with an outer membrane that is considered as one external layer. Inside the cytoplasm, we separate the osmotically active compartment containing small and medium channel-permeable osmolytes and larger channel-impermeable osmolytes from the osmotically inactive cytoplasmic compartment occupied by macromolecular species such as nucleic acids and proteins. Under steady growth conditions, the cytoplasmic membrane is pressed against the cell wall by turgor but has an extra area stored in small folds creating a surplus relative to the ‘visible’ area of the cell envelope. Upon osmotic downshift, the increased turgor pressure is predicted to expand the cell wall and completely unfold the inner membrane. It is only once the inner membrane is completely unfolded that the generated tension will be able to activate membrane-embedded MscS and MscL channels. Besides channels acting as discrete permeable elements, the membrane can form “cracks” that might open under very high (lytic) tensions. The cracks do not reseal immediately and thus may produce a protracted leakage. The cracks were modeled as large tension-gated channels with the energy of opening that is about twice of that for MscL and similar dimensions. Therefore, they open at tensions that are two times higher than the opening threshold for MscL. The total number of cracks and their rate of opening at the threshold, as well as their population heterogeneity, were allowed to vary (within 20-30%) and these additional adjustable parameters improved the fit. Except for the double-knockout mutant, the contribution of the cracks to the osmotic release is very small compared to the release through MS channels, meaning that the exact parameters of the cracks do not have a large effect on the scattering kinetics of the model; however, their activation reflects an excessive membrane tension and is a predictor of survival.

The model programmed as a set of cross-referenced ordinary differential equations to fit the time course of scattered light intensity explicitly takes into account the Rayleigh-Gans formalism including the effective radius and refractive index of cells, aerial excess of the inner membrane, elasticities of the membrane and the cell wall, water permeability of the membrane, starting concentrations, dependences of channel open probabilities on tension, osmolyte diffusion coefficients, and many others (see Supplement). The kinetic simulations were done for WT *V. cholerae* cells with both channels present, single Δ*mscL* and Δ*mscS* mutants, and the double channel knock-out (Figure 9). The curves are presented for one magnitude of shock (1200 ⟶ 300 mOsm) across the set of mutants. We should note that this is a strong downshift that kills about 70% of WT cells and wipes out Δ*mscS* and the double Δ*mscL*/Δ*mscS* knock-outs almost completely (Figure 3). Conspicuously, the Δ*mscL* mutant survives this shock better than WT.

**Figure 9.**
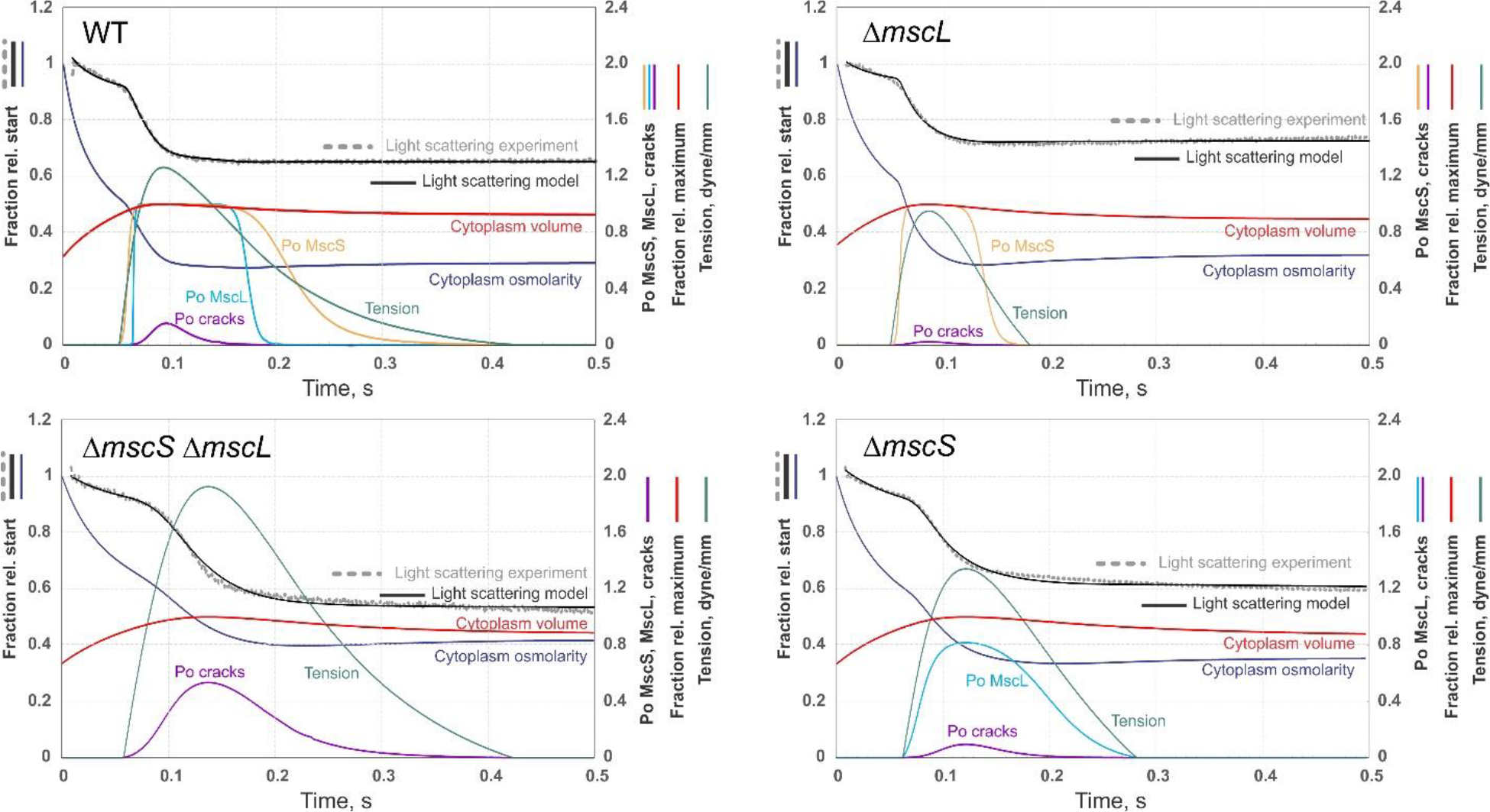
Kinetic models of *V. cholerae* WT and the three mutant’s responses to osmotic down-shock (from 1200 to 300 mOsm) that fit the experimental light scattering traces in Rayleigh-Gans approximation. The experimental traces are shown in gray, the fits in black and color coding of other model-deduced parameters is denoted by the curves. The legends by the left and right vertical scales specify the parameters represented in these scales. The modeling was done using COPASI (see the Supplement for details). On the panels, the curves belonging to the left vertical axis scale (cytoplasm osmolarity, measured and fitted light scattering) were normalized relative to the starting value. The right vertical axis specifies curves describing Po of MscL, MscS, and the cracks (by definition ranging between 0 and 1), as well as the volume of the cytoplasm expressed as a fraction (0 to 1) of the maximum observed value for the simulation. The right vertical axis additionally describes membrane tension expressed in dyne/mm to optimally fit the scale.

We see that the simulated scattering curve perfectly re-traces the experimental kinetics. The initial swelling reflected in approximately doubled cytoplasmic volume within the first ∼60 ms ends with activation of MscS (yellow curve) that is closely followed by a steep increase in Po for the MscL population (teal curve). After peaking at the relative level of about 1.3 dyne/mm (13 dyne/cm), tension (green curve, right y-axis) gradually decreases. During this process, MscL closes sharply, whereas MscS stays open for much longer. This sequential closing illustrates the role of low-threshold MscS in driving tension below the threshold for MscL, thus producing a fast and complete closure of MscL population. This important role of MscS in the process of quenching channel activity and membrane resealing after a near-lytic shock has been recently described for the *E. coli* MscS-MscL tandem (Moller et al., 2023). When tension exceeds the level of 1.2 dyne/mm, some number of cracks stochastically appear in the membrane which indicates that the shock of this magnitude is damaging even for WT (Figure 3).

The double mutant devoid of both channels experiences an extended swelling period (100 ms), during which tension approaches 1.95 dyne/mm causing a much higher probability of cracks, which correlates with a near-zero viability. The most viable Δ*mscL* mutant steeply opens its full MscS population at the earliest moment after the shock onset (∼50 ms), which reflects its low-tension activation threshold. It expediently releases osmolytes such that tension surge is curbed at 0.5 dyne/mm, resulting in minimal membrane damage with almost no cracks. The Δ*mscS* mutant, carrying the full complement of MscL channels, shows a delayed activation consistent with the MscL’s higher threshold, resulting in tension surging to 1.4 dyne/mm, which correlates with a low viability (Figure 3). The time to the onset of channel activation appears to be a critical factor for survival (see also comparison of experimental curves in Figure 6).

We should note that membrane cracks are the least defined physical entities and represent the most speculative aspect of the model. We introduced them to explain the time course of the double Δ*mscL*Δ*mscS* mutant response which points to the presence of some conductive/permeable structures appearing under very high tension. Besides crack formation, we expect that lesions in the cell wall occurring under extreme turgor can be another form of mechanical damage resulting in low survival. At this stage, we have no means to extract this type of outcome from the light scattering data, and other methods would be required.

## Discussion

The high adaptability of *Vibrio cholerae* to a variety of environments, from high-osmotic gut content to rainwater, facilitates its environmental stability and transmission, producing outbreaks. This osmotic stability is imparted by a set of tension-activated mechanosensitive channels that function as osmolyte release valves and quickly dissipate excessive osmotic gradients to rescue cells from lysis. The two major contributors to osmotic permeability are the low-threshold channel MscS and high-threshold MscL. When the knockout strains were first generated and functionally characterized in *Escherichia coli*, a wide held impression was that MscS and MscL were partially redundant and rescued shocked cells equally well (Levina et al., 1999). Subsequent studies have shown that the two channels are not redundant and work in tandem (Moller et al., 2023).

The engineered Δ*mscS*, Δ*mscL*, and the double Δ*mscS* Δ*mscL* strains from WT *V. cholerae* O395 reveal clear and correlated electrophysiological, release kinetic and survival phenotypes. Our data obtained in these strains indicate that MscS is the main contributor to the osmotic cell survival, whereas MscL alone poorly rescues cells and in the absence of MscS even becomes toxic. MscS activates earlier thereby reducing the time and extent of cell swelling. MscL activates later, upon tension, and when present alone, it leaves more time for water influx which damages the cell. The experimentally observed delayed kinetics of release in Δ*mscS* strains directly supports the notion of the competitive nature of osmotic rescuing when the efflux of osmolytes must match and outpace the water influx in order to curb the surge of tension. Kinetic models provide a quantitative estimation of the tension surge magnitude which clearly depends on the timing of channel activation by tension.

The analysis of stopped-flow light scattering traces clearly delineates kinetically separable phases of the permeability response (Figure 4) and reconstructs the events that take place *in vivo*. The sequence includes (1) initial cell swelling producing dilution of the cytoplasm and simultaneous distension of the peptidoglycan and outer membrane, (2) complete unfolding of the inner membrane generates tension which reaches the threshold for MS activation, (3) a fast osmolyte release phase through open channels, (4) the following process of water expulsion, and (5) slower volume/concentration adjustments possibly related to the recoil of the stretched peptidoglycan and resealing of the membrane. Kinetic modeling using COPASI provides a plausible quantitative picture which includes permeabilities for water and osmolytes, mechanical parameters of the cell (distensibility of the cell envelope and the inner membrane surplus) and presents the estimated dynamics of parameters ‘hidden’ from the view such as cell volume, cytoplasmic osmolarity, membrane tension, open probabilities for each of the channels and the kinetics of resealing. The application of this fast-mixing technique gives us precise answer to a simple question of how quickly does the release mechanism in *Vibrio cholerae* respond to a sudden medium dilution. Under extreme shocks, cells swell within 40 ms and release up to 17% of their dry weight within the next 20 ms. The best osmotic survival correlates with low extent of swelling (ES), short times to both release onset (tro) and steepest point (tsp) and, importantly, low downward slope at the end of the trace which signifies complete resealing of the membrane. These advantageous characteristics were found in strains expressing MscS.

In contrast to MscL which behaves as a two-state channel, MscS has the ability to inactivate from the closed state under moderate near-threshold tensions (Akitake et al., 2005; Cetiner et al., 2018). The inactivation of *V. cholerae* MscS has been reported previously (Rowe et al., 2013). When inactivated, MscS becomes tension insensitive. In *E. coli*, this distinctive property of MscS fulfills the special function of terminating the massive permeability response when tension drops down to MscS’s activation threshold. The mere presence of MscS at the end of the permeability response also silences the MscL channel by driving tension below its activation threshold. This is the picture that we now observe in *V. cholerae*. While the poorly surviving Δ*mscS* strain shows a nearly normal degree of swelling that should not inevitably lead to mechanical rupture of the cell, the kinetic traces demonstrate a continued release phase indicated by the negative slope due to the inability of the dominant MscL channel to close completely or inactivate. At the end of the active period, MscL likely remains in a low-open-probability state preventing complete membrane resealing (Figures 5 and 7). In the presence of active MscS, which lowers tension, pacifies MscL and then inactivates, the final levels in most traces are flat.

The mechanosensitive channel of small conductance (MscS) alone has a strong impact on survival possibly because it is also overexpressed in the Δ*mscL* strain suggesting a functional and transcriptional cross-talk between the two major osmolyte release valves (Figure 8). A similar effect of *mscL* deletion on the level of *mscS* mRNA was recently observed in *E. coli* (Moller et al., 2023).

We argue that the MscS channel that is capable of inactivation is a critical component of the release system allowing for proper termination of the hypoosmotic response. Moreover, if a Δ*mscL* mutant survives abrupt down-shocks even better than a strain with both channels, then why is MscL needed and preserved in evolution? We presume that the presence of a high-threshold non-inactivating channel such as MscL is therefore equally important. While the Δ*mscL* mutant demonstrates increased survival under single down-shock events, substantial inactivation of the MscS population under a graded shock scenario is possible. For instance, a prolonged exposure to a moderately hypoosmotic medium (pre-shock) would lead to massive inactivation of MscS, leaving *V. cholerae* susceptible to membrane damage under further stronger down-shocks. An exposure to non-lethal concentrations of antibiotics compromising the peptidoglycan layer and increasing its distensibility may produce tension and similarly inactivate MscS. The presence of the non-inactivating MscL population eliminates this risk.

We conclude that the osmotic rescuing in *Vibrio cholerae* is a kinetics-based mechanism where the two mechanosensitive channels, MscS and MscL, act in tandem. While the functional interaction between these two structurally unrelated species still needs to be explored in more detail, our data shows that MscS is the most important component of the system and that it can fulfill its rescuing function by itself. MscL is never found in any species alone because it requires MscS for proper termination of the osmotic permeability response.

## Acknowledgements

This research was supported by NIH R01AI135015 to SS. This research was also supported in part by a grant to the University of Maryland from the Howard Hughes Medical Institute through the Science Education Program to KR. This material is based upon work supported by the National Science Foundation Graduate Research Fellowship Program to EM under Grant No. DGE 1840340. Any opinions, findings, and conclusions or recommendations expressed in this material are those of the author(s) and do not necessarily reflect the views of the National Science Foundation.

We thank Anthony Schams for assistance with stopped-flow MATLAB fitting and analysis. We also thank Dr. Ugur Çetiner for his assistance in interpreting stopped-flow data.

