## Supplemental Material for "The dynamic hypoosmotic response of *Vibrio cholerae* relies on the mechanosensitive channel MscS"

BIORXIV/2023/539864 - Version 2
Supplemental material

### The dynamic hypoosmotic response of *Vibrio cholerae* relies on the mechanosensitive channel MscS

Kristen Ramsey[1,4], Madolyn Britt[1,2], Joseph Maramba[1], Blake Ushijima[5,6], Elissa Moller[1,2], Andriy Anishkin[1], Claudia Häse[6], and Sergei Sukharev[1,3]

[1]Department of Biology, [2]Biophysics Graduate Program, [3]Institute for Physical Science and Technology, University of Maryland, College Park, MD. [4]Department of Microbial Pathogenesis, Yale School of Medicine, New Haven, CT, [5]Biology and Marine Biology, University of North Carolina Wilmington, Wilmington, NC. [6]Carlson College of Veterinary Medicine, Oregon State University, Corvallis, OR.

RT-qPCR experiments

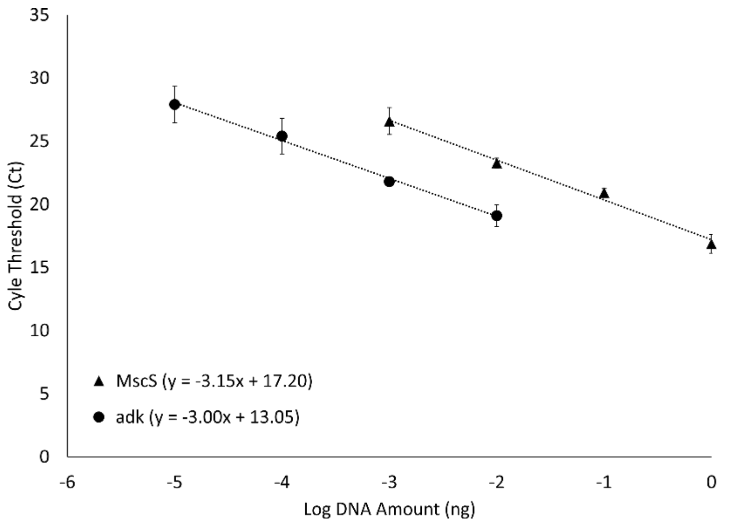


Figure S1. Primer efficiency plots for RT-qpCR experiments. The primer efficiency was 2.07 (105%) with a slope of -3.15 for the MscS primers and 2.14 (107%) with a slope of -3.00 for the adk primers.

RT-qPCR Primers

qPCR O395 MscS F: 5-GGCTCGGTTGATTCTATCCA-3
qPCR O395 MscS R: 5-GAAACCCCAATCACCATGTC-3
qPCR adk O395 F: 5-TCGATGCAGGACAATTGGTA-3
qPCR adk O395 R: 5-CAATCACATCATCCGCTACG-3

Simplification of the Rayleigh-Gans theory for kinetic fitting of scattering traces

The key dynamic variable that we need to fit to experimental data is the time course of the light scattered from the bacterial suspension. The directional intensity of the scattered light depends on multiple factors including the experimental setup, bacterial cell shape and the cytoplasmic content. For a suspension of micron-sized particles, the scattering is usually described in Rayleigh-Gans approximation 1.

$$I = \frac{8\pi^4 r^6 \eta_0^4}{R^2 \lambda^4} \cdot \frac{\left[\left(\frac{\eta}{\eta_0}\right)^2 - 1\right]^2}{\left[\left(\frac{\eta}{\eta_0}\right)^2 + 2\right]^2} \cdot \nu I_0 V [1 + cos^2(\theta)] \cdot P(\theta) \qquad (1)$$

where $I$ is the intensity of the scattered light at a given direction and distance; $r$ is the radius of the spherical equivalent of the actual particle (i.e. *E. coli* bacterium); $\eta_0$ is the index of refraction of the suspending medium; $\eta$ is the index of refraction of the particle; $V$ is the volume illuminated; $I_0$ is the intensity of the incident light: $\nu$ is the concentration of particles; $\theta$ is the angle of observation; $R$ is the distance of the observer from the sample; and $\lambda$ is the wavelength of light in vacuum. The factor $[1 + cos^2(\theta)]$, depends on the angle of observation, $\theta$, relative to the forward direction of the illuminating beam. For a given physical setup, most of these factors can be combined into an empirical constant. $P(\theta)$ is the appropriate correction function to compensate for the light interference within the particle.

To enable quantitative estimation of bacterial properties from the light scattering kinetics, the intensity equation can be simplified by separating parameters describing the experimental setup (scattering angle, distance, amplification, background light etc.), which usually remain constant throughout the experiment, from the dynamics occurring within the bacterial population, which include changes in the effective size (swelling/shrinkage) and the intracellular refractive index (Foladori et al., 2008; Koch, 1961; Koch et al., 1996). It was shown (Ball and Ramsden, 1998; Craig et al., 1995; Moller et al., 2023) that contributions of both small solutes and macromolecules to the refractive index scale linearly with their total mass fraction in the solution (dry weight). Because the release of the cytoplasmic content is dictated by the size cut-off of MS channels, all molecular species inside the cell can be divided in two categories – permeable (ions and small solutes which can cross the membrane through the channel) and impermeable (larger substances, proteins, nucleic acids etc.).

The refraction index of the bacterial cytoplasm approximated as a uniform solution, can therefore be presented as a linear combination:

$$\eta = \eta_w + \frac{d\eta}{dC_p} C_p + \frac{d\eta}{dC_i} C_i = \eta_w + \left(\frac{d\eta}{dC_p} m_p + \frac{d\eta}{dC_i} m_i\right) / V_c \qquad (2)$$

Where $\eta$ is the refraction index of the cytoplasm, $\eta_w$ is the refraction index of water, $C_p$ and $m_p$ are the concentration and total mass of the permeable species, and $C_i$ and $m_i$ are the respective parameters of the impermeable species, $V_c$ is the total volume of the bacterial cytoplasm.
By separating the instrumentation-related terms that into one constant *A* from time-dependent variables characterizing cell behavior, the equation for the intensity of the scattered light can be rewritten to:

$$I_r = I_b + Ar^6 \cdot \frac{\left[\left(\frac{\eta_w + 3\left(S_p m_p + S_i m_i\right)/4\pi r^3}{\eta_0}\right)^2 - 1\right]^2}{\left[\left(\frac{\eta_w + 3\left(S_p m_p + S_i m_i\right)/4\pi r^3}{\eta_0}\right)^2 + 2\right]^2} \qquad (3)$$

There $I_r$ is the recorded light intensity, $I_b$ is the background (stray) light component, $r$ the radius of the spherical equivalent of the bacterium, $\eta_w$ refraction index of water, $\eta_0$ refraction index of extracellular medium, $m_p$ and $m_i$ the masses of the permeable and impermeable solutes respectively, $S_p$ and $S_i$ the scaling coefficients between their concentrations and contributions to the refraction index of the medium ($S_p = d\eta/dC_p$ , $S_i = d\eta/dC_i$, see equation 2) and $A$ is the scaling coefficient for the amplitude of the scattered light that combines components for the instrumental amplification ($A_i$), system geometry, angular dependence, light intensity etc.

During the osmotic downshock, two of the variables in the equation 3 will undergo quick changes: the effective radius of the bacterium ($r$), and the mass of the permeable osmolytes within the cells ($m_p$). The rest of the terms in the equation can be considered near-constant in time. We expect that upon osmotic downshift bacteria will swell producing some effect of cytoplasm dilution, and then reach their maximum size when mechanosensitive channels open and the release phase begins. During the release phase we expect that the cell geometry remains relatively stable, and the changes in $I_r$ are now defined mostly by the remaining mass of permeable osmolytes inside the cell. During the release phase, $I_r$ becomes proportional to the square of the internal solute mass. In the case of moderate osmotic shrinking when osmolytes remain in the cell while water is exiting, the intensity of light scattering scales as $(1/r^2)$, i.e. inverse of the cell surface area, or as volume in the power of -2/3 (Koch, 1961; Koch et al., 1996).

Further simplification of equation (2) comes from the notion that the numerator is much smaller than the denominator, so the scaling of $I_r$ is quadratic with respect to the term $\left(S_p m_p + S_i m_i\right)$ , and in the course of osmolyte efflux, the ratio of refractive indexes between the cell and its environment becomes the major time-dependent parameter. With the assumption that contributions of permeable ($S_p$) and impermeable ($S_i$) osmolytes to the refractive index are the same (Moller et al., 2023) and the cell volume changes are small, the time course of the scattering signal can be presented as:

$$I = I_0 + S(m_i + m_p(t))^2 \qquad (4)$$

where $I$ is light intensity, $I_0$ is background light intensity, $S$ is the scaling coefficient, $m_i$ is the mass of impermeable osmolytes, and $m_p(t)$ is the mass of permeable osmolytes as a function of time $t$. Assuming exponential release of the permeable fraction of osmolytes, equation 4 can be presented in this form:

$$I = I_0 + S(m_i + m_{p0}e^{\frac{t_0 - t}{\tau}})^2 \qquad (5)$$

where $m_{p0}$ is the initial mass of the permeable osmolytes before the onset of release, and $t_0$ is the time at which the exponential release has started.

Details of kinetic simulations with COPASI

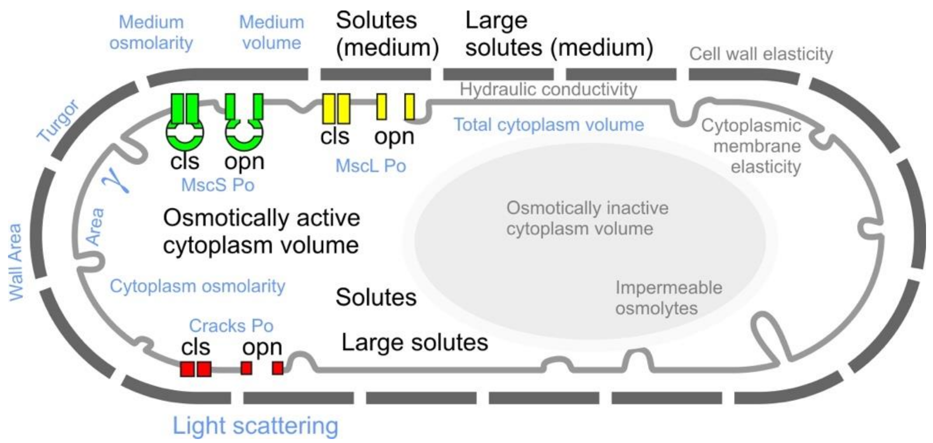


Figure S2. Key elements of the kinetic model of the bacterial response to osmotic shock. Independent variables are in black font, dependent values calculated based on those are in light blue, and static parameters are in light grey.

The bacterial osmotic response involves multiple interacting components with non-linear dependences and feedbacks, therefore the exact time course of the changes is difficult to predict analytically. However, it can be done by a numerical simulation given the known starting state and the relationships between the elements implemented as a system of differential equations. We have used COPASI, an advanced simulator for biochemical networks (Mendes et al., 2009), to model the dynamics of the bacterial response to osmotic shock. Our model (Figure S4) includes three compartments: (1) an osmotically active cytoplasm delineated by the cytoplasmic membrane containing impermeable osmolytes, as well as small and large permeable osmolytes, (2) an osmotically inactive cytoplasmic compartment occupied by macromolecular species such as DNA and proteins, and (3) the external medium which may contain small and large osmolytes. The cytoplasmic membrane embeds mechanosensitive channels MscS and/or MscL acting as permeable elements, and, besides the channels, the membrane can form “cracks” that might open under high tensions. The important mechanical component is the elastic cell wall (peptidoglycan) surrounding the cytoplasmic membrane. Under resting conditions when cells are grown under constant osmolarity, the membrane has a certain surplus of area stored in folds so that the surface area of the cell wall is slightly smaller than the area of fully unfolded cytoplasmic membrane.

The programmed sequence of events includes the following. With the onset of hypoosmotic shock, the external osmolarity drops and water starts flowing through the cytoplasmic membrane with the rate proportional to the hydraulic permeability and the osmolyte concentration gradient. As the osmotically active volume in the model increases, the turgor builds up, and the cell wall stretches in accordance to its rigidity (elasticity). The increasing turgor counters the water influx, and in the absence of osmolyte permeation, swelling would stop when the driving force from the concentration gradient is compensated by the hydrostatic pressure. Note that with the cell wall stretching and membrane unfolding, the cell volume increases and the internal osmolytes get diluted to some extent, which is tracked in our COPASI model and produces a dynamic adjustment of the water flow. Before this equilibrium state is reached, swelling may lead to a complete unfolding of the cytoplasmic membrane which starts stretching. As membrane tension increases, the open probability of the MscS and MscL channels rises. The process of channel activation is described in the model through rate equations based on experimental kinetic and thermodynamic data (Cetiner et al., 2018; Chiang et al., 2004; Sukharev et al., 1999). When channels (or membrane cracks) open, osmolytes get released with rates defined by their concentration gradients and permeability of a particular open channel for the given osmolyte type. Only small osmolytes can permeate through MscS, whereas both small and larger osmolytes can exit through MscL and the cracks). In the course of osmolyte release, the difference between the cytoplasm and medium osmolarities drops and the water flow ceases. Pressed by the recoil of the stretched cell wall, water leaves the cell and the cytoplasm volume decreases. With that, the tension in the cytoplasmic membrane goes down and channels gradually return to the closed state.

In order to describe the time courses of these coupled variables, the model needs parameters such as elasticity of the cell wall, aerial excess of the membrane, water permeability of the cytoplasmic membrane, starting concentrations, midpoints and slopes of channel open probabilities on tension, osmolyte diffusion coefficients, etc. Many of them are known from the literature and/or based on our own experimental measurements. Some require adjustment for a particular system. COPASI enables an automated search for parameters (within optional constraints) that optimize the fit between the model and multiple experimental measurements. We have used the light scattering of bacterial suspensions in the model to fine-tune some of the parameters (starting from the values found in the literature) to optimize the fit to the experimentally measured time course of the scattering triggered by abrupt osmotic shocks.

Below is the description of parameters used in simulations.

**Compartments (size dynamically changes with time):**

| Parameter | Comment |
|---|---|
| V_medium | Volume of the external medium |
| V_cytoplasm | Total cytoplasm volume |
| V_cytoplasm_OA | Osmotically active volume, derived from a total cell volume and a solid base volume of 41% |
| membrane_cyt | Cytoplasmic membrane |

**Variables (value dynamically changes with time):**

| Parameter | Comment |
|---|---|
| Crack_cls | Potential (closed) cracks in the cytoplasmic membrane |
| Crack_opn | Open cracks in the cytoplasmic membrane |
| LRG_cyt_d | Permeant osmolyte in the cytoplasmic core - needs time to diffuse to the surface |
| LRG_cyt_s | Permeant osmolyte concentration in the cytoplasm, near the cytoplasmic membrane - available for release |
| LRG_med | Permeant osmolyte concentration int he external medium |
| MscL_cls | Closed MscL |
| MscL_opn | Open MscL |
| MscS_cls | Closed MscS |
| MscS_inc | Inactivated MscS |

| | |
|---|---|
| MscS_opn | Open MscS |
| SLT_cyt_d | Permeant osmolyte in the cytoplasmic core - needs time to diffuse to the surface |
| SLT_cyt_s | Permeant osmolyte concentration in the cytoplasm, near the cytoplasmic membrane - available for release |
| SLT_med | Permeant osmolyte concentration in the external medium |

#### Calculated parameters (based on the changing compartments and variables):

| Parameter | Comment |
|---|---|
| Area_cell_wall | Total cell wall area |
| Area_membrane_cyt | Area of the cytoplasmic membrane. Starts to stretch when the cell wall area exceeds the unfolded cytoplasmic membrane |
| I | Intensity of the light scattering |
| Osm_cyt | Intracellular osmotically active concentration |
| Osm_med | Extracellular osmotically active concentration |
| Osm_med_shock_timecourse | Osmolarity change during the Osmotic shock. Starts at time t_shock_0 and has a certain mixing time t_mix |
| Po_Crack | Probability of a crack to be in an open state - calculated as a fraction of the open crack relative to the total crack |
| Po_MscL | Probability of MscL to be in an open state - calculated as a fraction of the open MscL relative to the total MscL |
| Po_MscS | Probability of MscS to be in an open state - calculated as a fraction of the open MscS relative to the total MscS |
| radius_cytoplasm | Effective radius of the bacterial cell (spherical approximation) |
| Tension | Lateral tension in cytoplasmic membrane, in N/m |
| Turgor | Turgor pressure [MPa] |
| VP | Non-turgid volume |

#### Common fixed parameters:

| Parameter | Comment |
|---|---|
| Ai | Scaling coefficient for the intensity of the light scattering |
| Area_membrane_cyt__unfolded | Unfolded area of the cytoplasmic membrane |
| Area_membrane_cyt__unfolded_f | Unfolded membrane area as a fraction of the starting area of the cell wall area |
| elasticity_membrane_cyt | Elasticity of the cytoplasmic membrane. Converts fractional expansion of the membrane (membrane expansion area over the relaxed area dA/A, unitless) into tension (N/m). Reference point is 10 dyne/cm (0.01 N/m) tension at 4% (0.04) expansion |
| Fc2p | Factor converting concentrations in M to pressures in MPa |
| Hydrvl_cond | Hydraulic conductivity of the cytoplasmic membrane |
| Ib | Intensity of the background light scattering |
| k | Boltzmann constant |
| mol | Mol 6.022*10^23 Mole number |
| n_cytoplasm | Refraction index of the cytoplasm |
| n_LB | Refraction index of LB medium |
| n_medium | Refraction coefficient of the external medium |
| n_water | Water refraction index |

| | |
|---|---|
| N2uM | Factor converting number of molecules in µM concentrations per cell |
| Osm_cyt_0 | Initial total cellular osmolyte concentration |
| Osm_cyt_imp_0 | Initial non-permeable cellular osmolyte concentration |
| Osm_med_0 | Initial osmolarity of the external medium, mOsm |
| Osm_med_shock | Change in the osmolyte concentration in the external medium, uM, for the osmotic stress. Can be positive (hyperosmotic) or negative (hypoosmotic) |
| R | Gas constant |
| rigidity | Cell wall rigidity |
| S_imp | Slope of the refraction index relative to the concentration of impermeable solutes/gel |
| S_LRG | Slope of the refraction index relative to the concentration of Mscs-permeable solutes |
| S_SLT | Slope of the refraction index relative to the concentration of Mscs-permeable solutes |
| T | Temperature in kelvin |
| t_mix | Mixing time [s] of the medium after the osmotic stress |
| t_shock_0 | Time [s] before the osmotic stress begins |
| Turgor_0 | Initial turgor pressure |
| Turgor2Osm | Conversion between turgor and osmolarity |
| V_cytoplasm_0 | Initial total cell volume |
| V_cytoplasm_imp | Solid/excluded/ minimal volume of the cell. Fixed value based on the initial fraction. |
| V_cytoplasm_imp_f | Minimal cell volume (as a fraction of total). Fixed starting value. |
| V_cytoplasm_OA_f | Osmotically Active cytoplasmic volume as a fraction of total cytoplasmic volume. Fixed starting value. |
| VP0 | Non-turgid volume - starting value |

#### Reactions and processes:

| **MscS_activation** | **Transition between the closed and open states of MscS** |
|---|---|
| **Parameter** | **Comment** |
| dAcb | Change in the expansion area between the closed state and the energy barrier |
| dAob | Change in the expansion area between the open state and the energy barrier |
| dE | Total energy cost of the reaction |
| dEb | The height of the energy barrier |
| h | Heterogeneity coefficient for the channel population (0 - completely heterogeneous, 1 - completely homogeneous) |
| Vtrans | The rate of transition |

| **SLT_permeation_MscS** | **Permeation of the small solutes through MscS** |
|---|---|
| **Parameter** | **Comment** |
| Po | Po of the channel |
| Vmax | The maximum rate |

| **SLT_diffusion** | **Diffusion of the small solutes (with delay kinetics)** |
|---|---|

| Parameter | Comment |
|---|---|
| a | Exponential coefficient for the diffusion equation |
| f | Referene raction for the diffusion equation |
| So | Reference concentration for the diffusion equation |
| Vmax | The maximum rate |

| **MscS_inactivation** | **Transition between the closed and inativated states of MscS** |
|---|---|
| **Parameter** | **Comment** |
| dAcb | Change in the expansion area between the closed state and the energy barrier |
| dAob | Change in the expansion area between the open state and the energy barrier |
| dE | Total energy cost of the reaction |
| dEb | The height of the energy barrier |
| h | Heterogeneity coefficient for the channel population (0 - completely heterogeneous, 1 - completely homogeneous) |
| Vtrans | The rate of transition |

| **Crack_opening** | **Transition between the closed and open states of the cracks in the membrane** |
|---|---|
| **Parameter** | **Comment** |
| dAcb | Change in the expansion area between the closed state and the energy barrier |
| dAob | Change in the expansion area between the inactivated state and the energy barrier |
| dE | Total energy cost of the reaction |
| dEb | The height of the energy barrier |
| h | Heterogeneity coefficient for the channel population (0 - completely heterogeneous, 1 - completely homogeneous) |
| Vtrans | The rate of transition |

| **LRG_permeation_Crack** | **Permeation of the large solutes through the cracks** |
|---|---|
| **Parameter** | **Comment** |
| Po | Po of the crack |
| Vmax | The maximum rate |

| **SLT_permeation_Crack** | **Permeation of the small solutes through the cracks** |
|---|---|
| **Parameter** | **Comment** |
| Po | Po of the crack |
| Vmax | The maximum rate |

*Unfolding of the membrane*

At rest, the cytoplasmic membrane has folds that take ~5% of the total area of the outer cell envelope which includes the cell wall and the outer membrane considered as one (observed cell surface). Under osmotic shock, the turgor gradually stretches the outer envelope; the area of the cytoplasmic membrane at the same time stays the same just until the envelope's area increases by 5%. After that, the area of the outer cell envelope becomes equal to the cytoplasmic membrane area, which now allows for tension generation:

$$\text{Area\_membrane\_cyt} = \begin{cases} \text{Area\_cell\_wall} > \text{Values[Area\_membrane\_cyt\_\_unfolded].InitialValue}, & \text{Area\_cell\_wall} \\ \text{else}, & \text{Values[Area\_membrane\_cyt\_\_unfolded].InitialValue} \end{cases}$$

The area of the cell wall was calculated based on the total cell cytoplasm volume:

$$\text{Area\_cell\_wall} = (36 \cdot \pi)^{\frac{1}{3}} \cdot V_{\text{V\_cytoplasm}}^{\frac{2}{3}}$$

The cytoplasmic volume is a sum of osmotically-active and -inactive volumes:

$$V_{\text{V\_cytoplasm}} = \text{Values[V\_cytoplasm\_imp].InitialValue} + V_{\text{V\_cytoplasm\_OA}}$$

The inactive volume is fixed. The osmotically-active volume is allowed to change due to the water flow in and out of the cell. The water flow (volume change) was proportional to the difference between the osmotic pressures (based on the difference of the solute osmolarities inside and outside of the bacterium) and turgor (based on the current cytoplasm volume and cell wall elasticity):

$$\frac{d V_{\text{V\_cytoplasm\_OA}}}{d t} = \text{Values[Hydrvl\_cond].InitialValue} \cdot \text{Area\_cell\_wall} \cdot (\text{Values[Fc2p].InitialValue} \cdot \text{Values[R].InitialValue} \cdot \text{Values[T].InitialValue} \cdot ([\text{Osm\_cyt}] - [\text{Osm\_med}]) - \text{Turgor})$$

*Elasticity of the cell wall*

The elasticity coefficient of the cell wall assumed to be constant. It is used as a coefficient in the equation defining the turgor pressure in response to the change in cytoplasm volume:

$$\text{Turgor} = \begin{cases} V_{\text{V\_cytoplasm}} > \text{VP}, & \text{Values[rigidity].InitialValue} \cdot \ln\left(\frac{V_{\text{V\_cytoplasm}}}{\text{VP}}\right) \\ \text{else}, & 0 \end{cases}$$

In this expression, the turgor pressure is proportional to cell wall rigidity, and logarithm of the relative change in volume of the cytoplasm normalized to the resting volume. The equation is based on the equation for the turgor response in yeast, as described by Schaber et al. (Schaber et al., 2010) which, in turn, was derived from the "linear elastic theory" saying that the change in turgor pressure P is proportional to a relative change in cell volume.

*Water efflux*

Water flow through the cytoplasmic membrane was supposed to be fast enough so that the osmotically-active cytoplasmic volume was set to adjust immediately to match the balance of the osmotic gradient and turgor pressure of the cell wall (same equation as above):

$$\frac{d V_{\text{V\_cytoplasm\_OA}}}{d t} = \text{Values[Hydrvl\_cond].InitialValue} \cdot \text{Area\_cell\_wall} \cdot (\text{Values[Fc2p].InitialValue} \cdot \text{Values[R].InitialValue} \cdot \text{Values[T].InitialValue} \cdot ([\text{Osm\_cyt}] - [\text{Osm\_med}]) - \text{Turgor})$$

The “hydraulic conductivity” for water is assumed to be constant (i.e. opening of MS channels does not increase the overall permeability presumably dominated by the lipid bilayer of the cytoplasmic membrane).

#### Equations:

$$\frac{\mathrm{d}V_{\text{V\_cytoplasm\_OA}}}{\mathrm{d}t} = \text{Values[Hydrvl\_cond].InitialValue}\cdot\text{Area\_cell\_wall}\cdot(\text{Values[Fc2p].InitialValue}\cdot\text{Values[R].InitialValue}\cdot\text{Values[T].InitialValue}\cdot([\text{Osm\_cyt}]-[\text{Osm\_med}])-\text{Turgor})$$

$$V_{\text{V\_medium}} = 11\cdot\text{Values[V\_cytoplasm\_0].InitialValue} - V_{\text{V\_cytoplasm}}$$

$$V_{\text{V\_cytoplasm}} = \text{Values[V\_cytoplasm\_imp].InitialValue} + V_{\text{V\_cytoplasm\_OA}}$$

$$\begin{aligned}\frac{\mathrm{d}([\text{SLT\_cyt\_s}]\cdot V_{\text{V\_cytoplasm\_OA}})}{\mathrm{d}t} = & -(\text{Vmax}_{(\text{SLT\_permeation\_MscS})}\cdot\text{Po\_MscS}\cdot([\text{SLT\_cyt\_s}]-[\text{SLT\_med}])) \\ & -(\text{Vmax}_{(\text{SLT\_permeation\_Crack})}\cdot\text{Po\_Crack}\cdot([\text{SLT\_cyt\_s}]-[\text{SLT\_med}])) \\ & +V_{\text{V\_cytoplasm\_OA}}\cdot\left(\frac{\text{Vmax}_{(\text{SLT\_diffusion})}\cdot([\text{SLT\_cyt\_d}]-[\text{SLT\_cyt\_s}])}{1+e^{a_{(\text{SLT\_diffusion})}\cdot\left(\frac{[\text{SLT\_cyt\_d}]}{\text{So}_{(\text{SLT\_diffusion})}}-f_{(\text{SLT\_diffusion})}\right)}}\right)\end{aligned}$$

$$[\text{Osm\_cyt}] = [\text{SLT\_cyt\_d}]+[\text{SLT\_cyt\_s}]+[\text{LRG\_cyt\_d}]+[\text{LRG\_cyt\_s}]+\frac{\text{Values[Osm\_cyt\_imp\_0].InitialValue}\cdot\text{Compartments[V\_cytoplasm\_OA].InitialVolume}}{V_{\text{V\_cytoplasm\_OA}}}$$

$$\begin{aligned}\frac{\mathrm{d}([\text{SLT\_med}]\cdot V_{\text{V\_medium}})}{\mathrm{d}t} = & +(\text{Vmax}_{(\text{SLT\_permeation\_MscS})}\cdot\text{Po\_MscS}\cdot([\text{SLT\_cyt\_s}]-[\text{SLT\_med}])) \\ & +(\text{Vmax}_{(\text{SLT\_permeation\_Crack})}\cdot\text{Po\_Crack}\cdot([\text{SLT\_cyt\_s}]-[\text{SLT\_med}]))\end{aligned}$$

$$[\text{Osm\_med}] = \text{Osm\_med\_0}+\text{Osm\_med\_shock\_timecourse}+[\text{SLT\_med}]+[\text{LRG\_med}]$$

$$\frac{\mathrm{d}([\text{MscS\_opn}]\cdot V_{\text{membrane\_cyt}})}{\mathrm{d}t} = +V_{\text{membrane\_cyt}}\cdot\left(\text{Vtrans}_{(\text{MscS\_activation})}\cdot\left(e^{\frac{h_{(\text{MscS\_activation})}\cdot(-(\text{dEb}_{(\text{MscS\_activation})}-\text{Tension}\cdot\text{dAob}_{(\text{MscS\_activation})}))}{k\cdot T}}\cdot[\text{MscS\_cls}]-e^{\frac{h_{(\text{MscS\_activation})}\cdot(-(\text{dEb}_{(\text{MscS\_activation})}-\text{dE}_{(\text{MscS\_activation})}-\text{Tension}\cdot\text{dAcb}_{(\text{MscS\_activation})}))}{k\cdot T}}\cdot[\text{MscS\_opn}]\right)\right)$$

$$\begin{aligned}\frac{\mathrm{d}([\text{MscS\_cls}]\cdot V_{\text{membrane\_cyt}})}{\mathrm{d}t} = & -V_{\text{membrane\_cyt}}\cdot\left(\text{Vtrans}_{(\text{MscS\_activation})}\cdot\left(e^{\frac{h_{(\text{MscS\_activation})}\cdot(-(\text{dEb}_{(\text{MscS\_activation})}-\text{Tension}\cdot\text{dAob}_{(\text{MscS\_activation})}))}{k\cdot T}}\cdot[\text{MscS\_cls}]-e^{\frac{h_{(\text{MscS\_activation})}\cdot(-(\text{dEb}_{(\text{MscS\_activation})}-\text{dE}_{(\text{MscS\_activation})}-\text{Tension}\cdot\text{dAcb}_{(\text{MscS\_activation})}))}{k\cdot T}}\cdot[\text{MscS\_opn}]\right)\right) \\ & -V_{\text{membrane\_cyt}}\cdot\left(\text{Vtrans}_{(\text{MscS\_inactivation})}\cdot\left(e^{\frac{h_{(\text{MscS\_inactivation})}\cdot(-(\text{dEb}_{(\text{MscS\_inactivation})}-\text{Tension}\cdot\text{dAob}_{(\text{MscS\_inactivation})}))}{k\cdot T}}\cdot[\text{MscS\_cls}]-e^{\frac{h_{(\text{MscS\_inactivation})}\cdot(-(\text{dEb}_{(\text{MscS\_inactivation})}-\text{dE}_{(\text{MscS\_inactivation})}-\text{Tension}\cdot\text{dAcb}_{(\text{MscS\_inactivation})}))}{k\cdot T}}\cdot[\text{MscS\_inc}]\right)\right)\end{aligned}$$

$$\frac{\mathrm{d}([\text{MscS\_inc}]\cdot V_{\text{membrane\_cyt}})}{\mathrm{d}t} = +V_{\text{membrane\_cyt}}\cdot\left(\text{Vtrans}_{(\text{MscS\_inactivation})}\cdot\left(e^{\frac{h_{(\text{MscS\_inactivation})}\cdot(-(\text{dEb}_{(\text{MscS\_inactivation})}-\text{Tension}\cdot\text{dAob}_{(\text{MscS\_inactivation})}))}{k\cdot T}}\cdot[\text{MscS\_cls}]-e^{\frac{h_{(\text{MscS\_inactivation})}\cdot(-(\text{dEb}_{(\text{MscS\_inactivation})}-\text{dE}_{(\text{MscS\_inactivation})}-\text{Tension}\cdot\text{dAcb}_{(\text{MscS\_inactivation})}))}{k\cdot T}}\cdot[\text{MscS\_inc}]\right)\right)$$

$$\begin{aligned}\frac{\mathrm{d}([\text{LRG\_cyt\_s}]\cdot V_{\text{V\_cytoplasm\_OA}})}{\mathrm{d}t} = & -(\text{Vmax}_{(\text{LRG\_permeation\_MscL})}\cdot\text{Po\_MscL}\cdot([\text{LRG\_cyt\_s}]-[\text{LRG\_med}])) \\ & +V_{\text{V\_cytoplasm\_OA}}\cdot\left(\frac{\text{Vmax}_{(\text{LRG\_diffusion})}\cdot([\text{LRG\_cyt\_d}]-[\text{LRG\_cyt\_s}])}{1+e^{a_{(\text{LRG\_diffusion})}\cdot\left(\frac{[\text{LRG\_cyt\_d}]}{\text{So}_{(\text{LRG\_diffusion})}}-f_{(\text{LRG\_diffusion})}\right)}}\right) \\ & -(\text{Vmax}_{(\text{LRG\_permeation\_Crack})}\cdot\text{Po\_Crack}\cdot([\text{LRG\_cyt\_s}]-[\text{LRG\_med}]))\end{aligned}$$

$$\frac{\mathrm{d}([\text{LRG\_med}]\cdot V_{\text{V\_medium}})}{\mathrm{d}t} = +(\text{Vmax}_{(\text{LRG\_permeation\_Crack})}\cdot\text{Po\_Crack}\cdot([\text{LRG\_cyt\_s}]-[\text{LRG\_med}]))$$

$$\frac{d([\text{Crack\_cls}]\cdot V_{\text{membrane\_cyt}})}{dt} = -V_{\text{membrane\_cyt}}\cdot\left(\text{Vtrans}_{(\text{Crack\_opening})}\cdot\left(e^{\frac{h_{(\text{Crack\_opening})}\cdot(-(\text{dEb}_{(\text{Crack\_opening})}-\text{Tension}\cdot\text{dAob}_{(\text{Crack\_opening})}))}{k\cdot T}}\cdot[\text{Crack\_cls}]-e^{\frac{h_{(\text{Crack\_opening})}\cdot(-(\text{dEb}_{(\text{Crack\_opening})}-\text{dE}_{(\text{Crack\_opening})}-\text{Tension}\cdot\text{dAcb}_{(\text{Crack\_opening})}))}{k\cdot T}}\cdot[\text{Crack\_opn}]\right)\right)$$

$$\frac{d([\text{Crack\_opn}]\cdot V_{\text{membrane\_cyt}})}{dt} = +V_{\text{membrane\_cyt}}\cdot\left(\text{Vtrans}_{(\text{Crack\_opening})}\cdot\left(e^{\frac{h_{(\text{Crack\_opening})}\cdot(-(\text{dEb}_{(\text{Crack\_opening})}-\text{Tension}\cdot\text{dAob}_{(\text{Crack\_opening})}))}{k\cdot T}}\cdot[\text{Crack\_cls}]-e^{\frac{h_{(\text{Crack\_opening})}\cdot(-(\text{dEb}_{(\text{Crack\_opening})}-\text{dE}_{(\text{Crack\_opening})}-\text{Tension}\cdot\text{dAcb}_{(\text{Crack\_opening})}))}{k\cdot T}}\cdot[\text{Crack\_opn}]\right)\right)$$

$$\text{V\_cytoplasm\_OA\_f} = 1-\text{Values[V\_cytoplasm\_imp\_f].InitialValue}$$

$$\text{V\_cytoplasm\_imp} = \text{Values[V\_cytoplasm\_0].InitialValue}\cdot\text{Values[V\_cytoplasm\_imp\_f].InitialValue}$$

$$\text{VP0} = \text{Compartments[V\_cytoplasm].InitialVolume}\cdot e^{\frac{-(\text{Values[Turgor\_0].InitialValue})}{\text{Values[rigidity].InitialValue}}}$$

$$\text{N2uM} = \frac{1e21}{\text{Values[mol].InitialValue}\cdot\text{Compartments[V\_cytoplasm\_OA].InitialVolume}}$$

$$\text{Osm\_cyt\_0} = \text{Values[Osm\_cyt\_imp\_0].InitialValue}+[\text{SLT\_cyt\_d}]\text{\_0}+[\text{SLT\_cyt\_s}]\text{\_0}+[\text{LRG\_cyt\_d}]\text{\_0}+[\text{LRG\_cyt\_s}]\text{\_0}$$

$$\text{Turgor} = \begin{cases} V_{\text{V\_cytoplasm}} > \text{VP}, & \text{Values[rigidity].InitialValue}\cdot\ln\left(\frac{V_{\text{V\_cytoplasm}}}{\text{VP}}\right) \\ \text{else}, & 0 \end{cases}$$

$$\text{Po\_MscS} = \frac{[\text{MscS\_opn}]}{[\text{MscS\_opn}]+[\text{MscS\_cls}]+[\text{MscS\_inc}]}$$

$$\text{Turgor2Osm} = \frac{\text{Turgor}}{\text{Values[R].InitialValue}\cdot\text{Values[T].InitialValue}\cdot\text{Values[Fc2p].InitialValue}}$$

$$\text{VP} = \text{Compartments[V\_cytoplasm].InitialVolume}\cdot\left(\frac{\text{Compartments[V\_cytoplasm].InitialVolume}}{\text{VP0}}\right)^{\frac{-(\text{Values[rigidity].InitialValue})}{\text{rigidity}}}$$

$$\text{Area\_cell\_wall} = (36\cdot\pi)^{\frac{1}{3}}\cdot V_{\text{V\_cytoplasm}}^{\frac{2}{3}}$$

$$\text{Osm\_med\_shock\_timecourse} = \begin{cases} \text{Time} < \text{t\_shock\_0}, & 0 \\ \text{else}, & \text{Osm\_med\_shock}\cdot\left(1-e^{\frac{-(\text{Time}-\text{t\_shock\_0})}{\text{t\_mix}}}\right) \end{cases}$$

$$\text{Area\_membrane\_cyt} = \begin{cases} \text{Area\_cell\_wall} > \text{Values[Area\_membrane\_cyt\_\_unfolded].InitialValue}, & \text{Area\_cell\_wall} \\ \text{else}, & \text{Values[Area\_membrane\_cyt\_\_unfolded].InitialValue} \end{cases}$$

$$\text{Tension} = \frac{\text{Values[elasticity\_membrane\_cyt].InitialValue}\cdot(\text{Area\_membrane\_cyt}-\text{Values[Area\_membrane\_cyt\_\_unfolded].InitialValue})}{\text{Values[Area\_membrane\_cyt\_\_unfolded].InitialValue}}$$

$$\text{Po\_Crack} = \frac{[\text{Crack\_opn}]}{[\text{Crack\_opn}]+[\text{Crack\_cls}]}$$

$$\text{radius\_cytoplasm} = \left(\frac{3}{4\cdot\pi}\cdot V_{\text{V\_cytoplasm}}\right)^{\frac{1}{3}}$$

$$\text{n\_medium} = \frac{\text{Values[n\_LB].InitialValue}\cdot(\text{Values[Osm\_med\_0].InitialValue}+\text{Osm\_med\_shock\_timecourse})+\text{Values[n\_water].InitialValue}\cdot(-(\text{Osm\_med\_shock\_timecourse}))}{\text{Values[Osm\_med\_0].InitialValue}}$$

$$\text{n\_cytoplasm} = \text{Values[n\_water].InitialValue}+\frac{\text{Values[S\_imp].InitialValue}\cdot\text{Values[Osm\_cyt\_imp\_0].InitialValue}\cdot\text{Compartments[V\_cytoplasm].InitialVolume}}{V_{\text{V\_cytoplasm}}}+\text{Values[S\_SLT].InitialValue}\cdot([\text{SLT\_cyt\_d}]+[\text{SLT\_cyt\_s}])+\text{Values[S\_LRG].InitialValue}\cdot([\text{LRG\_cyt\_d}]+[\text{LRG\_cyt\_s}])$$

$$I = \text{Values[Ib].InitialValue}+\text{Values[Ai].InitialValue}\cdot\text{radius\_cytoplasm}^{6}\cdot\left(\frac{\left(\frac{\text{n\_cytoplasm}}{\text{n\_medium}}\right)^{2}-1}{\left(\frac{\text{n\_cytoplasm}}{\text{n\_medium}}\right)^{2}+2}\right)^{2}$$

Statistical treatment of osmotic viability assays

| Relative to WT | 800 | 600 | 400 | 350 | 300 | 250 | 200 | 150 |
|---|---|---|---|---|---|---|---|---|
| Δ*mscS* | 4.2E-3* | 2.8E-6* | 1.5E-7* | 1.3E-3* | 3.6E-3* | 2.5E-2* | 2.9E-2* | 4.3E-2* |
| Δ*mscL* | 2.3E-1 | 2.4E-1 | 5.6E-7# | 2.7E-3# | 1.1E-3# | 6.9E-5# | 6.6E-2 | 7.5E-1 |
| Δ*mscS* Δ*mscL* | 1.4E-4* | 2.3E-6* | 4.9E-8* | 1.1E-3* | 3.6E-3* | 2.2E-2* | 2.8E-2* | 4.3E-2* |

Table S1. **P-values for t-tested osmotic viability assays.** P-values from two-tail two-sample t-tests assuming unequal variances used to determine statistical significance between osmotic viability experiments. The strains listed in the first column were all analyzed for statistically significant differences relative to WT. Significant values are below 0.05 and are marked with an asterisk for values lower than WT or a pound sign for values that are higher than WT.

Data and statistics for figure 7

| Fold Volume Change | 900 | 650 | 500 | 400 | 250 | 100 |
|---|---|---|---|---|---|---|
| WT | 2.0 ± 0.3 | 2.1 ± 0.1 | 2.3 ± 0.0 | 2.3 ± 0.1 | 2.2 ± 0.2 | 2.0 ± 0.1 |
| ΔmscS | 1.4 ± 0.4 | 2.5 ± 0.2* | 2.4 ± 0.3 | 2.4 ± 0.1 | 2.4 ± 0.2 | 2.2 ± 0.1* |
| ΔmscL | 1.9 ± 0.3 | 2.1 ± 0.2 | 2.2 ± 0.2 | 2.0 ± 0.1* | 2.2 ± 0.1 | 2.1 ± 0.2 |
| ΔmscSΔmscL | 2.2 ± 0.2 | 2.5 ± 0.2* | 2.4 ± 0.1* | 2.8 ± 0.5 | 2.7 ± 0.4 | 2.6 ± 0.3* |

Table S2. **Fold volume change values for WT and mutants.** The value of the fold volume change in response to rapid dilution in stopped flow experiments expressed as the mean ± standard deviation calculated from N trials. N=4 (WT), N=4 (ΔmscS), N=5 (ΔmscL), N=4 (ΔmscSΔmscL).

| Fold Volume Change | 900 | 650 | 500 | 400 | 250 | 100 |
|---|---|---|---|---|---|---|
| WT | | | | | | |
| ΔmscS | 0.129 | *0.025* | 0.432 | 0.364 | 0.242 | *0.028* |
| ΔmscL | 0.959 | 0.665 | 0.131 | *0.007* | 0.861 | 0.362 |
| ΔmscSΔmscL | 0.296 | *0.022* | *0.046* | 0.167 | 0.061 | *0.025* |

Table S3. **P-values for t-tested fold volume change.** P-values from two-tail two-sample t-tests assuming unequal variances used to determine statistical significance for the fold volume change as determined from stopped flow experiments. The strains listed in the first column were all analyzed for statistically significant differences relative to WT. Significant values are below 0.05 and italicized.

| Time to Release Onset | 900 | 650 | 500 | 400 | 250 | 100 |
|---|---|---|---|---|---|---|
| WT | 1.1E-1 ± 4.5E-2 | 8.9E-2 ± 1.4E-2 | 6.8E-2 ± 2.6E-3 | 5.8E-2 ± 2.1E-3 | 5.1E-2 ± 9.5E-4 | 4.3E-2 ± 2.9E-4 |
| ΔmscS | 1.3E-1 | 1.3E-1 ± 1.6E-2* | 8.9E-2 ± 4.7E-3* | 7.5E-2 ± 1.8E-3* | 6.3-2 ± 5.0E-4* | 5.4E-2 ± 2.9E-3* |
| ΔmscL | 1.3E-1 ± 1.9E-2 | 9.1E-2 ± 4.4E-3 | 7.0E-2 ± 2.1E-3 | 5.5E-2 ± 7.5E-4* | 5.1E-2 ± 1.3E-3 | 4.4E-2 ± 2.6E-3 |
| ΔmscSΔmscL | 1.7E-1 ± 1.3E-2 | 1.2E-1 ± 1.6E-2 | 8.7E-2 ± 4.8E-3* | 8.8E-2 ± 1.3E-2* | 7.1E-2 ± 1.2E-2* | 6.1E-2 ± 1.0E-2* |

Table S4. **Time to release onset values for WT and mutants.** The value of the time to release onset in response to rapid dilution in stopped flow experiments expressed as the mean ± standard deviation calculated from N trials. N=4 (WT), N=4 (ΔmscS), N=5 (ΔmscL), N=4 (ΔmscSΔmscL).

| Time to Release Onset | 900 | 650 | 500 | 400 | 250 | 100 |
|---|---|---|---|---|---|---|
| WT | | | | | | |
| ΔmscS | | *0.021* | *5.6E-4* | *1.6E-5* | *3.4E-6* | *0.004* |
| ΔmscL | 0.433 | 0.790 | 0.232 | *0.043* | 0.761 | 0.597 |
| ΔmscSΔmscL | 0.072 | 0.067 | *0.009* | *0.020* | *0.041* | *0.040* |

Table S5. **P-values for t-tested time to release onset.** P-values from two-tail two-sample t-tests assuming unequal variances used to determine statistical significance for the time to release onset as determined from stopped flow experiments. The strains listed in the first column were all analyzed for statistically significant differences relative to WT. Significant values are below 0.05 and italicized.

| Time to Steepest Point | 900 | 650 | 500 | 400 | 250 | 100 |
|---|---|---|---|---|---|---|
| WT | 2.1E-2 ± 8.4E-3 | 1.0E-1 ± 1.1E-2 | 8.1E-2 ± 2.1E-3 | 7.2E-2 ± 2.9E-4 | 6.4E-2 ± 1.2E-3 | 5.5E-2 ± 6.5E-4 |
| ΔmscS | 1.2E-2 ± 3.6E-3 | 1.6E-1 ± 2.1E-2* | 1.1E-1 ± 4.9E-3* | 9.4E-2 ± 4.2E-3* | 7.8E-2 ± 5.8E-4* | 6.8E-2 ± 4.8E-3* |
| ΔmscL | 1.0E-2 ± 9.5E-3 | 1.1E-1 ± 1.0E-3 | 8.2E-2 ± 7.6E-4 | 7.2E-1 ± 1.5E-3 | 6.7E-2 ± 2.1E-3 | 5.6E-2 ± 2.5E-3 |
| ΔmscSΔmscL | 2.3E-1 ± 3.4E-2* | 1.4E-1 ± 3.3E-3* | 1.2E-1 ± 2.3E-2* | 1.1E-1 ± 1.6E-2* | 8.6E-2 ± 1.3E-2* | 7.3E-2 ± 1.0E-2* |

Table S6. **Time to steepest point values for WT and mutants.** The value of the time to steepest point in response to rapid dilution in stopped flow experiments expressed as the mean ± standard deviation calculated from N trials. N=4 (WT), N=4 (ΔmscS), N=5 (ΔmscL), N=4 (ΔmscSΔmscL).

| Time to Steepest Point | 900 | 650 | 500 | 400 | 250 | 100 |
|---|---|---|---|---|---|---|
| WT | | | | | | |
| ΔmscS | 0.193 | *0.011* | *3.6E-4* | *0.002* | *3.5E-5* | *0.012* |
| ΔmscL | 0.155 | 0.213 | 0.561 | 1 | 0.070 | 0.544 |
| ΔmscSΔmscL | *0.009* | *0.003* | *0.036* | *0.021* | *0.043* | *0.040* |

Table S7. **P-values for t-tested time to steepest point.** P-values from two-tail two-sample t-tests assuming unequal variances used to determine statistical significance for the time to steepest point as determined from stopped flow experiments. The strains listed in the first column were all analyzed for statistically significant differences relative to WT. Significant values are below 0.05 and italicized.

| Fraction of Permeable Osmolytes | 900 | 650 | 500 | 400 | 250 | 100 |
|---|---|---|---|---|---|---|
| WT | 0.01 ± 0.00 | 0.04 ± 0.01 | 0.07 ± 0.01 | 0.11 ± 0.01 | 0.14 ± 0.00 | 0.17 ± 0.00 |
| ΔmscS | 0.03 ± 0.02 | 0.07 ± 0.02* | 0.11 ± 0.01* | 0.12 ± 0.01 | 0.13 ± 0.01 | 0.15 ± 0.01* |
| ΔmscL | 0.01 ± 0.00 | 0.03 ± 0.00 | 0.06 ± 0.00 | 0.07 ± 0.02* | 0.14 ± 0.01 | 0.16 ± 0.01 |
| ΔmscSΔmscL | 0.02 ± 0.01 | 0.05 ± 0.02 | 0.11 ± 0.02* | 0.13 ± 0.00 | 0.14 ± 0.00* | 0.16 ± 0.00* |

Table S8. **Fraction of permeable osmolytes values for WT and mutants.** The value of the fraction of permeable osmolytes in response to rapid dilution in stopped flow experiments expressed as the mean ± standard deviation calculated from N trials. N=4 (WT), N=4 (ΔmscS), N=5 (ΔmscL), N=4 (ΔmscSΔmscL).

| Fraction of Permeable Osmolytes | 900 | 650 | 500 | 400 | 250 | 100 |
|---|---|---|---|---|---|---|
| WT | | | | | | |
| ΔmscS | 0.159 | *0.017* | *6.2E-4* | 0.349 | 0.137 | *0.021* |
| ΔmscL | 0.391 | 0.166 | 0.198 | *0.003* | 0.729 | 0.712 |
| ΔmscSΔmscL | 0.127 | 0.319 | *0.024* | 0.079 | *0.005* | *0.015* |

Table S9. **P-values for t-tested fraction of permeable osmolytes.** P-values from two-tail two-sample t-tests assuming unequal variances used to determine statistical significance for the fraction of permeable osmolytes as determined from stopped flow experiments. The strains listed in the first column were all analyzed for statistically significant differences relative to WT. Significant values are below 0.05 and italicized.

| 1/Tau | 900 | 650 | 500 | 400 | 250 | 100 |
|---|---|---|---|---|---|---|
| WT | 23.4 ± 3.9 | 41.3 ± 5.5 | 66.2 ± 5.6 | 53.9 ± 0.5 | 68.3 ± 4.1 | 76.3 ± 1.3 |
| ΔmscS | 23.7 ± 10.3 | 23.2 ± 3.7* | 26.5 ± 0.7* | 41.3 ± 4.6* | 64.4 ± 2.7 | 63.2 ± 11.2 |
| ΔmscL | 13.1 ± 6.7* | 42.4 ± 1.7 | 68.9 ± 6.5 | 51.5 ± 1.8 | 58.7 ± 2.7* | 70.1 ± 1.4* |
| ΔmscSΔmscL | 8.7 ± 1.2* | 11.1 ± 0.8* | 15.1 ± 1.6* | 37.6 ± 11.5 | 56.1 ± 12.7 | 67.6 ± 7.0 |

Table S10. **1/Tau values for WT and mutants.** The value of the rate of release in response to rapid dilution in stopped flow experiments expressed as the mean ± standard deviation calculated from N trials. N=4 (WT), N=4 (ΔmscS), N=5 (ΔmscL), N=4 (ΔmscSΔmscL).

| 1/Tau | 900 | 650 | 500 | 400 | 250 | 100 |
|---|---|---|---|---|---|---|
| WT | | | | | | |
| ΔmscS | 0.957 | *0.016* | *7.9E-4* | *0.013* | 0.192 | 0.102 |
| ΔmscL | *0.024* | 0.761 | 0.521 | 0.062 | *0.012* | *0.005* |
| ΔmscSΔmscL | *0.002* | *0.011* | *6.6E-5* | 0.067 | 0.143 | 0.091 |

Table S11. **P-values for t-tested 1/tau.** P-values from two-tail two-sample t-tests assuming unequal variances used to determine statistical significance for the rate of osmolyte release as determined from stopped flow experiments. The strains listed in the first column were all analyzed for statistically significant differences relative to WT. Significant values are below 0.05 and italicized.

| Slope for the Final 100 ms | 900 | 650 | 500 | 400 | 250 | 100 |
|---|---|---|---|---|---|---|
| WT | 1.7E-2 ± 8.8E-3 | 6.5E-3 ± 6.0E-3 | 1.5E-2 ± 6.9E-3 | -4.4E-3 ± 5.0E-3 | -3.7E-2 ± 3.3E-3 | -3.5E-2 ± 9.1E-3 |
| ΔmscS | -2.6E-3 ± 1.1E-2* | 8.2E-3 ± 7.1E-3 | -1.4E-2 ± 1.3E-3* | -3.2E-2 ± 7.3E-3* | -4.2E-2 ±5.7E-3 | -4.7E-2 ± 4.9E-3 |
| ΔmscL | 1.4E-2 ± 1.1E-2 | 1.7E-2 ± 3.7E-3 | 1.5E-2 ± 9.6E-3 | 2.4E-2 ± 7.8E-3* | -3.7E-2 ± 1.3E-2 | -3.8E-2 ± 1.0E-2 |
| ΔmscSΔmscL | 1.3E-2 ± 7.8E-3 | 1.4E-2 ± 1.9E-2 | -1.4E-2 ± 3.7E-2 | -3.5E-2 ± 3.1E-2 | -4.1E-2 ± 9.7E-3 | -4.6E-2 ± 4.5E-3 |

Table S12 **Slope values for WT and mutants.** The value of the slope for the final 100 ms of release in response to rapid dilution in stopped flow experiments expressed as the mean ± standard deviation calculated from N trials. N=4 (WT), N=4 (ΔmscS), N=5 (ΔmscL), N=4 (ΔmscSΔmscL).

| Slope for the Final 100 ms | 900 | 650 | 500 | 400 | 250 | 100 |
|---|---|---|---|---|---|---|
| WT | | | | | | |
| ΔmscS | *0.031* | 0.748 | *0.004* | *0.002* | 0.225 | 0.065 |
| ΔmscL | 0.709 | 0.082 | 0.991 | *7.8E-4* | 0.986 | 0.673 |
| ΔmscSΔmscL | 0.595 | 0.487 | 0.216 | 0.145 | 0.486 | 0.092 |

Table S13. **P-values for t-tested slope.** P-values from two-tail two-sample t-tests assuming unequal variances used to determine statistical significance for the slope of the final 100 ms of release as determined from stopped flow experiments. The strains listed in the first column were all analyzed for statistically significant differences relative to WT. Significant values are below 0.05 and italicized.
